# The HSP90–CDC37 Chaperone System Orchestrates RAF1 Kinase Activation Through a Pre-Dimerization Mechanism

**DOI:** 10.64898/2026.03.25.713956

**Authors:** Gonzalo Aizpurua, Pablo Mesa, Laura de-la-Puente-Ovejero, J. Rafael Ciges-Tomas, Lucia Lomba-Riego, Carmen G. Lechuga, Eduardo Zarzuela, Marta Isasa, Leander van der Hoeven, Jesper V. Olsen, Mariano Barbacid, Sara García-Alonso, Guillermo Montoya

**Affiliations:** Experimental Oncology Group, Tumor Biology Programme, Centro Nacional de Investigaciones Oncológicas (CNIO), Melchor Fernández Almagro 3, 28029 Madrid, Spain; Structural Molecular Biology Group, Protein Structure and Function Program, Novo Nordisk Foundation Centre for Protein Research, Department of Cellular and Molecular Medicine, Faculty of Health and Medical Sciences University of Copenhagen, Blegdamsvej 3B, Copenhagen, 2200, Denmark; Centro de Investigación Biomédica en Red de Cáncer (CIBERONC), Instituto de Salud Carlos III, 28029, Madrid, Spain; Proteomics Core Unit, Biotechnology Programme, Centro Nacional de Investigaciones Oncológicas (CNIO), Melchor Fernández Almagro 3, 28029 Madrid, Spain; Proteomics Program, Novo Nordisk Foundation Centre for Protein Research, Department of Cellular and Molecular Medicine, Faculty of Health and Medical Sciences University of Copenhagen, Blegdamsvej 3B, Copenhagen, 2200, Denmark

## Abstract

RAF kinases activate MEK in the RAS–MAPK signaling pathway, and changes in RAF kinase signaling have been linked to tumor formation. RAF1 requires the HSP90-CDC37 chaperone system for proper activation, but how the HSP90–CDC37 chaperone system regulates RAF kinase maturation remains enigmatic. We present novel cryo-EM structures of previously uncharacterized RAF1 chaperone complexes, including a 2:2:2 RAF1–HSP90–CDC37 complex (RRHCC), intermediate assemblies (RHCC), and a RAF1–HSP90–CDC37–p23 complex (RHCp23). These reveal an asymmetric stepwise folding mechanism unique among HSP90 kinase clients in which one RAF1 threads through the HSP90 lumen while another is captured in a “casting mold” formed by CDC37 and HSP90 that stabilizes the partially folded αC helix of RAF1. The RHCp23 structure shows how p23 cooperates with CDC37 to regulate ATP hydrolysis and client release. The HSP90-CDC37 system supports pre-dimerization of RAF1 and BRAF^V600E^ homodimers and RAF1 heterodimers, a mechanism unique to RAF among kinase clients of HSP90. Phosphoproteomics reveals selective activating phosphorylations within RRHCC. These RAF isoform complexes differentially activate MEK signaling and cell proliferation, establishing HSP90–CDC37 as not just a passive stabilizer but an active regulator of RAF signaling with therapeutic implications.

## INTRODUCTION

RAF kinases are central regulators of the mitogen-activated protein kinase (MAPK) signaling cascade^1,2^, which is critical for cell proliferation, differentiation, survival, and apoptosis^1^. In vertebrates, the RAF family comprises three isoforms—ARAF, BRAF and RAF1 (also known as CRAF)—which^2^ share a modular organization: an N-terminal regulatory region (CR1) containing the RAS-binding domain (RBD) and cysteine-rich domain (CRD); a central CR2 segment enriched for regulatory phosphorylation sites; and a C-terminal kinase domain (CR3)^2^. By phosphorylating their sole substrate, MEK1/2, RAF kinases play a key role in transducing extracellular signals to downstream effectors, ultimately leading to the activation of the extracellular signal-regulated kinases (ERK1/2)^1,2^. Despite their similar structures, the RAF kinases exhibit distinct biochemical properties, and their dysregulation by missense mutations^3^ or fusion protein formation^4,5^ results in oncogenic isoforms responsible for the onset of multiple tumor types.

The physiological regulation of RAF kinases involves a complex sequence of steps: recruitment to the membrane by RAS-GTP^6^ to increase their effective concentration^7,8^, maturation of certain unstable isoforms by HSP90-CDC37^9,10^, phosphorylation events that relieve inhibitory constraints or promote activation^11,12^, and finally dimerization^13,14^, which is stabilized by 14-3-3 binding to phosphorylated C-terminal sites and drives catalytic competence^12,15^. These processes are tightly regulated and provide different scenarios for an eventual therapeutic intervention^16^ ^17^. However, the mechanisms by which RAF activation is regulated during the transition from an immature/unstable kinase to a mature, dimerization-competent enzyme remain poorly understood.

RAF1 plays pivotal roles in signaling^10,18^, and is stabilized by the HSP90-CDC37 chaperone system, which assists its maturation into a functional conformation that is capable of activation and dimerization. While HSP90-CDC37 interact weakly with wild-type BRAF^10^, the oncogenic mutants of BRAF^9^, ARAF^10^ and RAF1^19^, are well-established clients of this chaperone system^9,20–22^. HSP90-CDC37 prevents RAF1 degradation and facilitates its dephosphorylation and phosphorylation^20^. The association of RAF1 with HSP90-CDC37 is found in cytosolic and membrane cellular fractions^23^, and is crucial for RAF1 activity and MAPK pathway signaling^19^. Once folded, RAF1 associates with 14-3-3 dimers in a phosphorylation dependent maner^24^, facilitating the formation of back-to-back homo-and heterodimers with other RAF family members^12,15^. Assembly with 14-3-3 dimers shifts RAF kinases into the catalytically competent conformations^25^ critical for MAPK signaling^13,14,26^. The association of certain RAF isoforms with HSP90-CDC37 suggests that the chaperone system may not only stabilize its RAF clients^24,27–29^ but shape the conformational and post-translational state that determines whether they can assemble into specific homo-and heterodimers, thereby modulating signaling outputs^12,15,29–31^. How the association of HSP90-CDC37 with certain RAF family isoforms controls their maturation and influences signaling output is unclear.

Here, we present the structure of previously uncharacterized intermediate assemblies of RAF1-chaperone complexes, including <u>R</u>AF1-<u>H</u>SP90-<u>C</u>DC37 complexes with 2:2:2 (RRHCC), 1:2:1 (RHC), and 1:2:2 (RHCC) stoichiometries, as well as the association of the RHC with the p23 cochaperone (RHCp23). These structures provide a complete landscape on how these chaperones cooperate to assist RAF1 folding, revealing key snapshots of the maturation process. The folding of the RAF clients of HSP90-CDC37 is specific, distinguishing RAF kinases from canonical clients like CDK4^32,33^. Furthermore, chaperone-assisted RAF dimers modulate MAPK pathway activity differentially, depending on the dimer composition. Collectively, these findings reveal the conformational landscape of RAF1 maturation, showing that RAF clients of HSP90-CDC37 undergo a unique pre-dimerization process. This regulatory mechanism goes beyond passive stabilization to integrate control of ATP hydrolysis, residue phosphorylation, and dimer readiness, providing a novel understanding of the role of chaperone systems in kinase regulation and signaling. These findings have implications for designing therapeutic strategies to disrupt oncogenic RAF signaling in cancer.

## RESULTS

### Structural heterogeneity of RAF1-chaperone complexes revealed by cryo-EM

To understand the maturation and activation of RAF1, we co-expressed RAF1 and CDC37 in Expi293F cells together with the KRAS^G12V^ mutant, which stabilizes the GTP bound state favoring the interaction with RAF1 RBD^34,35^. This strategy was followed to capture transient intermediates of KRAS with the kinase and the chaperone complex, which are challenging to reconstitute with recombinant proteins. The RAF1-chaperone complexes were isolated through two sequential rapid affinity purification steps targeting epitope tags first on RAF1 and then on KRAS (STAR Methods). The purified preparation was validated by SDS-PAGE, western blotting, and mass spectrometry, which confirmed the presence of HSP90, RAF1, KRAS, and CDC37 and post-translational phosphorylation of RAF1, HSP90, and CDC37 (Fig. S1, Table S1). The preparation was divided into two fractions: one was used directly for cryo-EM and phosphorylation analysis, and the remainder was subjected to size-exclusion chromatography for further biochemical characterization (Fig. S1C-D).

Cryo-EM analysis of vitrified samples revealed substantial compositional and conformational heterogeneity in RAF1-chaperone complexes (Fig. S2, Table S2). Despite biochemical evidence confirming KRAS association with the purified complexes (Fig. S1A-B), we did not observe any density corresponding to KRAS bound to the RAF1 RBD in any of the resolved structures. This absence suggests that KRAS engages RAF1 RBD through a highly flexible or transient interaction that precludes its visualization in the maps. Next, we detail how these structural snapshots provide unprecedented insight into previously unknown assembly intermediates during RAF1 maturation, revealing a stepwise chaperone-mediated folding mechanism.

### Architectures of the RAF1 chaperone complexes

The RRHCC complex unveils a novel asymmetric assembly in which two RAF1-CDC37 heterodimers associate with opposite faces of a closed HSP90 dimer (Fig. 1A–C, Fig. S3). The cryo-EM maps achieved sufficient resolution to enable atomic modeling of the complete HSP90 dimer with bound nucleotides (Fig. S3A-B), one full-length CDC37 protomer, a portion of the second CDC37 molecule, and distinct segments of both RAF1 CR3 kinase domains. Both CDC37 molecules engage their respective RAF1 protomers and dock onto the HSP90 molecular clamp on opposite sides of the dimer. This interaction is mediated by phosphorylation at serine 13 (pS13) on CDC37, consistent with our mass spectrometry data showing phosphorylation at this site (Table S1, Fig. S3C).

**Figure 1.**
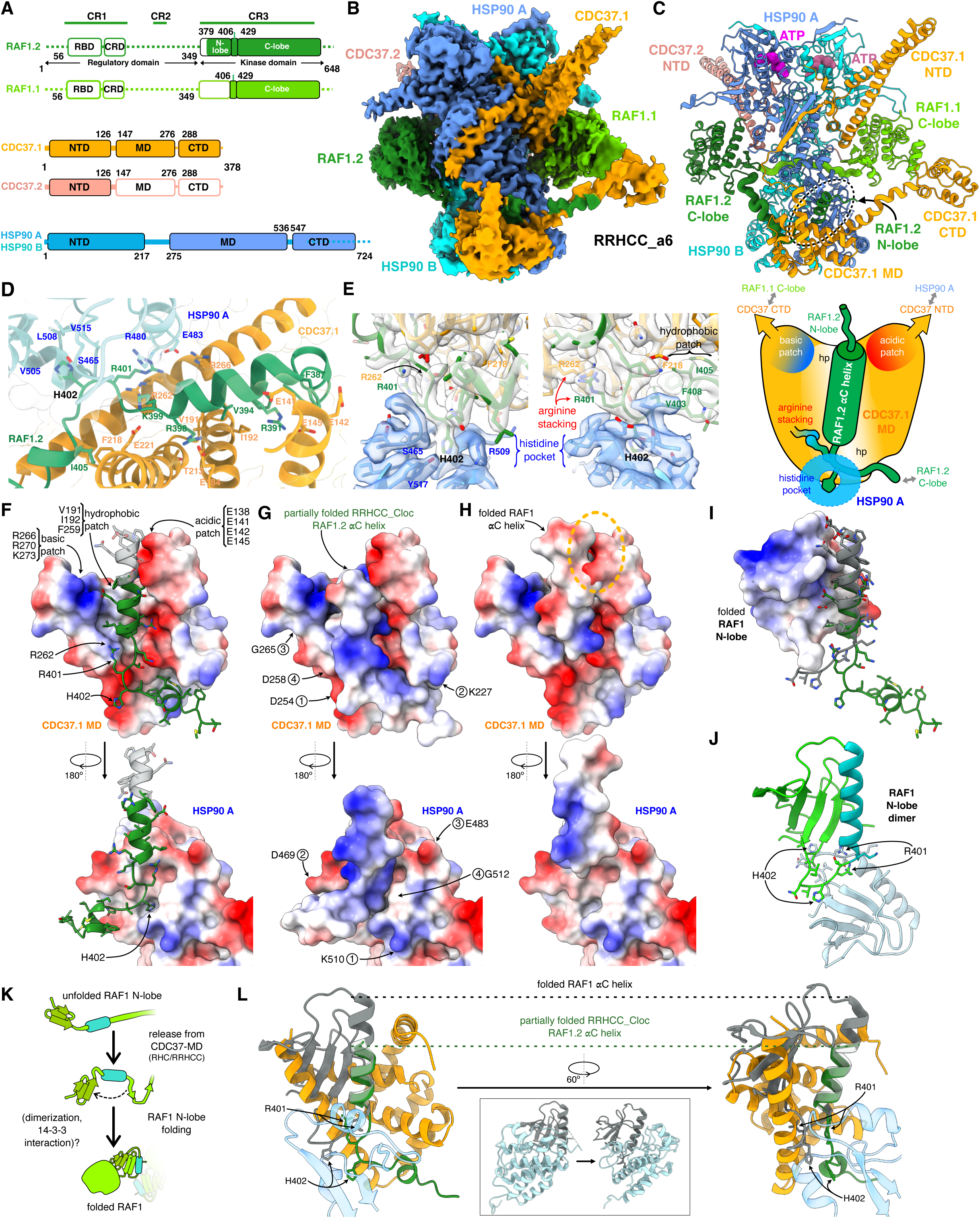
Cryo-EM structure of the RRHCC complex. **(A)** Schematic diagram showing the domain organization of the proteins present in the RRHCC complex. The dashed and non-colored sections indicate regions where there is no interpretable density. **(B)** Cryo-EM map of the RRHCC complex (RRHCC_a6) at 2.6 Å global resolution. The map is colored according to each subunit as in (A). **(C)** Cartoon representation of the RRHCC assembly in complex with ATP (sphere representation). The N-lobe sections of RAF1.1 and RAF1.2 associated with HSP90-CDC37 are depicted as thicker coils. The black oval highlights the interaction of RAF1.2 N-lobe with CDC37.1-MD and HSP90-A. **(D)** Zoom of the N-lobe of RAF1.2 showing an almost completely folded αC helix, which is captured between the CTD of HSP90-A and CDC37.1-MD. **(E)** Two perpendicular views of the interface formed by the RAF1.2 αC helix, CDC37.1-MD, and HSP90-A (green, orange, and blue, respectively) in the local map RRHCC_Cloc. Key interacting residues are labeled. The accompanying cartoon of αC in the mold illustrates the arrangement. The residues at the base of the αC helix set its position and register by engaging distinct hydrophobic (hp), basic, and acidic patches. **(F)** Surface representation of CDC37.1-MD (model from RRHCC_Cloc map), colored by electrostatic potential. Two opposite views are shown: CDC37.1-MD region (upper panel) and the corresponding HSP90-A interface (lower panel). RAF1.2 (green ribbon) fits into the central groove of CDC37.1 MD, which features two oppositely charged patches at its exit (upper panel). RAF1.2 residue H402 inserts into a pocket of HSP90-A (lower panel). A fully extended αC helix from the folded RAF1 structure (PDB: 8CPD, transparent grey) would clash with the acidic patch of CDC37.1-MD. Relevant RAF1.2 residues are labeled. **(G)** Same as in (F) with the αC of RAF1.2 displayed with its electrostatic potential surface, showing its complementation with the electrostatic potential of CDC37 and HSP90. Pairs of interacting residues between CDC37.1-MD and HSP90-A are indicated. **(H)** The αC helix of RAF1.2 is substituted with its fully folded conformation from PDB entry 8CPD to illustrate that the folded helix is incompatible with the acidic patch located at the exit of CDC37.1-MD. **(I)** Electrostatic potential surface of the folded RAF1 N-lobe, with the αC helix shown as a grey ribbon. For comparison, the fitted RAF1.2 model from the RRHCC_Cloc map is displayed in green in the same orientation as in (A). **(J)** Interface between the RAF1 N-lobes in a dimer (PDB:8CPD), with residues that contact CDC37-MD highlighted. One subunit is colored according to the scheme used in Fig. 1A. **(K)** Schematic illustrating how the secondary structure elements of the RAF1 N-lobe assemble into the final folded state. When the αC helix is released from the CDC37-MD, the segment of RAF1 that is clamped by HSP90 must be released upon HSP90 opening. Then, together with the final 30 N-terminal residues in the RRHCC, the kinase can fold the final two β-sheets. These β-sheets are inserted between the αC helix and the remaining β-strands building the RAF1 N-lobe. **(L)** Comparison of the model from the local map and the folded RAF1 N-lobe (PDB: 8CPD), aligned at the αC helix region. The inset shows the fully folded RAF1 structure, with the N-lobe in grey and the C-lobe in white. Also see Figure S2, SI1-4, and Table S2.

The RHC complex identified in this preparation closely resembles the structure we reported previously^24^. However, the improved data quality enabled exhaustive 3D classification focused on the middle domain of CDC37 (CDC37-MD), revealing the conformational landscape of this assembly (Fig. S2, S4, SI1-3) and allowing visualization of distinct ATP species within the HSP90 nucleotide-binding pockets (Fig. S4E). Notably, while some reconstructions show intact ATP molecules in both HSP90 protomers (RHC_c2, c4 classes), other classes reveal asymmetric nucleotide occupancy. Specifically, the HSP90 protomer directly interacting with the RAF1-CDC37 module contains ADP at its nucleotide-binding site (Site A), while the opposite HSP90 protomer retains intact ATP (Site B) (RHC_c1 and RHC_c3 classes) (Fig. S4E). These differences in nucleotide hydrolysis states correlate with the conformational variability observed in the CDC37-MD across different structural classes (Fig. S4D-E), suggesting that ATP hydrolysis at Site A is coupled to conformational changes in CDC37 that may regulate RAF1 loading or release.

**Figure 2.**
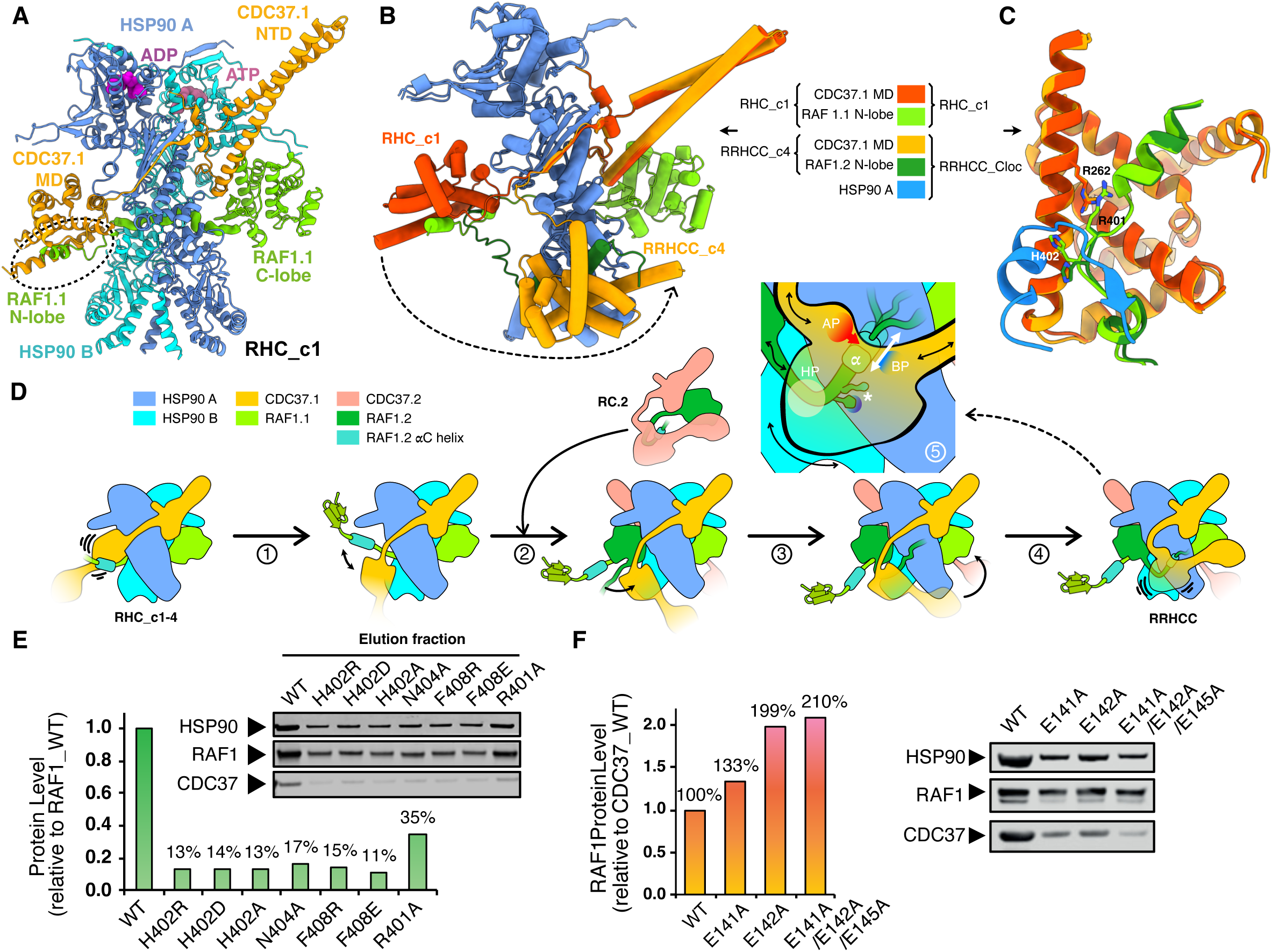
Interaction of RAF1.2. α**C in the mold. (A)** Ribbon representation of the RHC complex (RHC_c1). The black oval highlights the interaction between the partially folded RAF1.1 N-lobe and CDC37.1-MD in RHC, equivalent to the RAF1.2–CDC37.1-MD contact seen in RRHCC (Fig. 1C). **(B)** Simplified depiction of the proposed conformational change (black arrow) of the CDC37.1-MD between RHC_c1 and RRHCC_c4, suggesting the release of the αC helix of RAF1.1 and the capture of the very same element of RAF1.2. Only the latter allows the additional interaction with HSP90-A. Models were superimposed on their HSP90-A and RAF1.2 C-lobe. HSP90 B and CDC37.2 are hidden for clarity. **(C)** Comparison of the αC-helix interaction with CDC37.1-MD in the RHC_c1 (RAF1.1) and RRHCC_Cloc maps (RAF1.2). **(D)** Model of RRHCC assembly from a pre-formed RHC complex. The interaction between the RAF1.1 N-lobe and CDC37.1-MD is flexible (Fig. S4), but eventually the RAF1.1 αC helix fully folds and is released. Release of CDC37.1-MD then allows recruitment of an additional CDC37:RAF1 complex (2). The new RAF1.2 N-lobe binds the now-available CDC37.1-MD (3), together with HSP90-A, forming the casting mold in which the RAF1.2 αC helix becomes partially folded and transiently trapped (4). This assembly is continuously under strain due to its connections to distant parts of the complex, e.g. RAF1.1 C-lobe via CDC37.1-CTD, HSP90-A via CDC37-NTD and RAF1.2 C-lobe via its unfolded linker, which collectively challenge the stability and extension of the captured RAF1.2 αC helix (5). Also see Figures S4 and S7. **(E**) Immunoblot analysis of WT and mutant RAF1 proteins co-expressed with CDC37 and HT-KRAS^G12V^ in Expi293F cells following affinity purification.*Bottom.* Quantitative MS-based proteomics (iBAQ) using RAF1 common peptides to determine relative protein levels. Data are normalized to RAF1 WT levels. **(F)** *Right.* Immunoblot analysis of WT and mutant CDC37-St proteins co-expressed with RAF1-V5 and HT-KRAS^G12V^ in Expi293F cells following StrepTag affinity purification. *Left.* Quantitative MS-based proteomics (iBAQ) using RAF1 common peptides to determine relative protein levels. Data represent the mean ± SD of three independent biological replicates, normalized to CDC37 WT.

In addition, we identified the RHCC complex. In this assembly, which contains two protomers of CDC37, the conformation of RAF1 resembles that in the RHC complex^24^ (Fig. S5A-B). However, both HSP90 nucleotide-binding pockets are occupied by ADP (Fig. S5C). Interestingly, one CDC37 molecule lacks clearly associated RAF1 density, and only its N-terminal segment comprising the antiparallel β-sheet and the interface with HSP90-MD, including phosphorylation at S13, is well-defined (residues 2-133) (Fig. S5D-E). The absence of RAF1 kinase domain density associated with this second CDC37 molecule likely reflects a weaker interaction of RAF1 with HSP90, resulting in increased local flexibility that limits structural resolution in this region. This observation suggests that RHCC may represent either an early assembly intermediate where only one RAF1 molecule has been loaded or a late-stage complex where one RAF1 molecule has been released following maturation.

Finally, an extensive 3D classification and variability analysis of a second preparation enabled us to identify an additional particle population among RAF1 chaperone complexes leading to an RHC like complex with the p23 cochaperone bound (Fig. S2, SI3, Table S2, STAR Methods). The structure displays the C-lobe of RAF1 associated with HSP90 and CDC37, as in the RHC^24^, and a p23 protomer on the opposite side of the HSP90 closed clamp, representing an unique structure of a HSP90 in complex with a kinase client and two co-chaperones (see Structure of RHCp23 complex section).

### RAF1 exhibits distinct folding states in the RRHCC complex

The RRHCC complex captures two RAF1 molecules, each associated with its own CDC37 protomer, in strikingly different conformational states. The two RAF1 kinase domains share structural similarity in their C-lobes (Fig. 1A-C, Fig. S6), which superimpose closely with the corresponding region in the RAF1-14-3-3 active complex structure (1.0 Å RMSD for 151 Cα atoms, PDB: 8CPD)^30^. The separation of the C-and N-lobes hinders the folding of the kinase domain and the internal assembly of the Regulatory (R-spine) and Catalytic (C-spine) spines, which align the enzyme for efficient phosphotransfer. Consistent with previous observations^24,33^, the N-terminal domains (NTDs) of CDC37s mimic interactions normally formed between the N-and C-lobes of folded kinases, thereby stabilizing these separated subdomains during the maturation process. However, the two RAF1 protomers adopt radically different conformations within the HSP90-CDC37 assembly (Fig. S3A), breaking the inherent symmetry of the HSP90 dimer and revealing that the chaperone machinery employs distinct molecular strategies to capture and stabilize RAF1 at different stages of folding.

The first RAF1 protomer (RAF1.1), bound to CDC37.1 on one side of the HSP90 clamp, closely resembles the kinase conformation we previously observed in the RHC complex^24^. The interaction between the RAF1.1 C-lobe and CDC37.1 middle domain (CDC37.1-MD) mirrors that seen in RHC (Fig. 1B-C, Fig. S3A, Fig. S6A-B). In this configuration, the N-and C-lobes of RAF1.1 remain separated, with an elongated polypeptide segment spanning residues T413 to G426 of the N-lobe extending into the HSP90 lumen (Fig. S3D). This represents a canonical chaperone-client interaction where the kinase domain is held in an open immature state, displaying a folded C-lobe while the N-lobe, comprising the phosphate binding loop (P-loop), the αC and the N-terminal β−sheet, is unfolded and almost completely non-observed in the cryo-EM maps^24,28,33^.

The second RAF1 protomer (RAF1.2) adopts a strikingly different conformation on the opposite face of the HSP90 dimer, where the RAF1.2 C-lobe associates with CDC37.2 (Fig. 1A-C, Fig. S3A, S6A-B,). While RAF1.2 also displays separated N-and C-lobes, the extended Y411–L429 segment of the N-lobe is completely detached from the folded RAF1.2 C-lobe, stretching along the complex toward a novel interaction surface formed by CDC37.1-MD and the middle domain of HSP90 subunit A (HSP90-A-MD). Thus, RAF1.2 interacts with both CDC37 protomers. The remaining portion of the N-lobe (residues G410-F387) threads directly into this interface (Fig. 1C-D, Fig. S3A, S6C). Approximately 30 residues comprising β-strands 1–3 and the P-loop of the N-lobe are completely disordered and invisible in the density map of RAF1.2 (Fig. S6B-C). This extensive disorder suggests that these critical elements of the kinase ATP-binding pocket have not yet folded. However, the αC helix, a key regulatory element of kinase activity that is not visible in RAF1.1, is almost completely assembled in RAF1.2. This helix nestles snugly into a groove formed between HSP90-A-MD and the CDC37.1-MD, with H402 of RAF1.2 serving as an “anchor” that stabilizes the base of the αC helix (Fig. 1D-E). The groove acts as a molecular “casting mold” that cradles and stabilizes the partially folded αC helix, encasing it in two complementary surfaces (Fig. 1E-H). CDC37.1-MD provides a hydrophobic surface that accommodates the amphipathic character of the αC helix, mimicking aspects of its native environment in the folded kinase. However, whereas the αC helix is solvent-exposed in mature RAF1 monomers and dimers (Fig. 1I-L), in this intermediate, the opposite face of the helix contacts an adjacent acidic patch on CDC37. This contact appears to serve as a length-control mechanism, preventing full extension and premature folding of this secondary structure element (Fig. 1E-I, 1L).

### The histidine anchor locks the ***α***C helix in the mold

The H402 “anchor” of RAF1.2 is a conserved residue that is inserted into a specific pocket on the HSP90 surface (Fig. 1D-H, Fig. S3E). The neighboring residue, R401, forms a stacking interaction with R262 on CDC37.1-MD, further stabilizing this critical junction (Fig. 1E-F). The insertion of H402 into the HSP90 pocket is facilitated by a turn in the polypeptide chain created by a hydrogen bond between N404 and the backbone amide of L406. Together with hydrophobic interactions at the base of the helix, this histidine anchor, which in active kinases is positioned proximal to the R-spine^36^, creates a stable foundation that allows the αC helix to fold partially while remaining captured within the mold (Fig. 1L, Fig. S7). Interestingly, the H402 and the neighboring residues contribute to the dimer interface in the active kinase complex with 14-3-3^30^, while it is locked in the RRHCC.

### Conformational heterogeneity in the mold region

The mold region exhibits substantial conformational heterogeneity, giving rise to multiple structural classes of both RHC and RRHCC complexes (Fig. S2). CDC37.1-MD and CTD undergo major conformational changes that modulate formation of the αC helix in RAF1.2 and consequently influence N-lobe folding (Fig. S7). To address this intrinsic heterogeneity, we calculated a locally refined map (RRHCC_Cloc) and performed extensive 3D classification based on CDC37.1-MD variability (classes RRHCC_c1–c6) and CDC37.1-CTD engagement with the RAF1.1 C-lobe (classes RRHCC_a1–a6) (Fig. S2, Table S2, STAR Methods). This analysis revealed multiple conformational states that differ primarily in the orientation and extension of the αC helix, reflecting rigid-body movements of CDC37.1-MD relative to HSP90 (Figs. S8A–C, Fig. S7). These movements appear driven by the need to accommodate shifting protein-protein interfaces rather than motion along a single defined trajectory. The conformational flexibility of CDC37.1-MD is further constrained by multiple factors, including the polypeptide segment connecting the RAF1.2 C-lobe to the αC helix, the flexible linker between CDC37.1-MD and CDC37.1-NTD, and additional interactions between CDC37.1-CTD and the RAF1.1 C-lobe in a region that normally serves as the MEK-binding interface^30,31^ (Fig. 1B-C, Fig. S8D-F). These interactions, absent in the RHC, are strategically positioning the CDC37-CTD and may serve to prevent premature or inappropriate binding of immature RAF1 to its substrate MEK during the maturation process.

### Dynamic remodeling to trap a new kinase asssembles the mold

Despite the dramatically different overall conformations of RAF1.2 in the RRHCC compared to RAF1 in the RHC, the association of the αC helix with CDC37-MD is remarkably conserved, differing mainly in the orientation of the critical H402 residue (Fig. 2A-C). In the RRHCC complex, H402 of RAF1.2 inserts into the cleft on HSP90-A-MD, forming the histidine anchor (Fig. 1E). In contrast, in the RHC complex, the corresponding H402 is exposed to solvent (Fig. 2C). Nevertheless, the surrounding hydrophobic residues of the kinase and the stacked arginine residues occupy similar positions in both structures. This structural comparison suggests a dynamic mechanism where the association of a second RAF1-CDC37 client module to form the RRHCC complex induces a movement of CDC37.1-MD, which releases the αC helix of RAF1.1, and simultaneously captures and stabilizes the corresponding element of RAF1.2 through its new interaction with HSP90 (Fig. 2B-D, S8G). This model allows us to propose a loading pathway where a pre-assembled RAF1-CDC37 complex can be captured by HSP90 to form either the RHC complex or, upon binding of a second RAF1-CDC37 module, progress to the RRHCC state (Fig. 2D, S8H). Collectively, our structural analysis reveals that H402 constitutes a critical molecular node linking enzyme maturation with subsequent activation following binding to 14-3-3 (Fig. 1J-K). This dual role positions H402 as a critical integration point in RAF1 regulation, linking chaperone-mediated folding with activation mechanisms required for downstream signaling. The dynamic movements observed in the mold region challenge the stability of the RAF1.2 αC helix, preventing its full extension while maintaining a primed conformation that can eventually nucleate complete folding. This conformational checkpoint allows controlled dissociation from the chaperone machinery and final maturation of the N-lobe to produce a fully folded, catalytically competent RAF1 kinase.

### Mutations in the mold affect complex formation

To investigate the functional importance of the observed molecular interactions and test our model, we generated RAF1 and CDC37 mutants targeting key residues and isolated complexes from Expi293F cells. The relative abundance of RAF1 within these complexes was quantified by mass spectrometry and compared to wild-type protein levels (STAR Methods). Mutation of critical residues at the histidine anchor interface between RAF1 and CDC37 dramatically reduced RAF1 levels in isolated complexes by 65%–89% (Fig. 2E). This substantial reduction suggests that disrupting the histidine anchor leads to RAF1 destabilization and subsequent proteasomal degradation, consistent with previous observations that misfolded kinases are targeted for degradation^1,9^. In contrast, when the histidine anchor was preserved but the acidic patch on CDC37 was mutated to hydrophobic residues, we observed increased RAF1 levels in the complexes (Fig. 2F). This finding indicates that replacing the acidic residues facilitates complete folding of the αC helix and promotes RAF1 release from the chaperone machinery. The hydrophobic substitutions likely remove both the steric hindrance and the electrostatic repulsion that the acidic patch exerts on the helix, a disruptive effect that the single basic residue R391 can only partially counteract (Fig. 1F-I). These mutagenesis results underscore the functional importance of both the histidine anchor and the acidic patch as critical regulatory features controlling RAF1 maturation. These elements drive the dynamic conformational changes of the HSP90-CDC37 machinery observed in our structures and precisely modulate assembly of the αC helix within the casting mold (Fig. S7).

### Phosphorylation analysis of the RRHCC complex

Phosphorylation is a key mechanism controlling RAF kinase activity, dimerization, subcellular localization, and interaction with scaffold proteins^2,37^. Multiple phosphorylation sites, both activating and inhibitory, finely tune RAF function in response to upstream cues. Because the RRHCC complex was observed after co-expression with KRAS, we hypothesized that the phosphorylation status of RAF1 may differ in the RRHCC sample when compared to the RHC complex (STAR Methods). Hence, we performed a phosphoproteomics study to analyze the phosphorylation pattern of the sample used in the cryo-EM, where these subcomplexes have been observed. The samples were analyzed using a data-dependent acquisition (DDA) mass spectrometry (MS) experiment, with phosphorylation set as a variable modification in MaxQuant search (Fig. S1F-H). Phosphorylation sites were identified in all the complex subunits with a localization probability score greater than 75%, indicating confident assignment to a specific amino acid (Table S1, STAR Methods). We found several previously unreported phosphosites in RAF1, but we focused our analysis on the phosphorylation status of 16 well-characterized key residues^37^ (Fig. S1F). When compared with the previous RHC phosphoproteomics analysis^24^, nine sites (S29, S338, S339, S471, T491, S494, S497, S499 and S642) exhibited differential phosphorylation between the RHC and RRHCC complexes. These include both known inhibitory sites (S29, S642) and activating sites (S338, S339, S471, T491, S494, S497, S499)^38,39^, indicating a phosphorylation remodeling of RAF1 in the RRHCC complex. Notably, the inhibitory residues were not phosphorylated in the RRHCC complex, whereas the activating residues were specifically phosphorylated in the RRHCC complex but not in the RHC complex (Table S1, Fig. S1H). These activating residues are often associated with conformational changes that promote RAF dimerization and MAPK pathway signaling. This selective phosphorylation pattern suggests that RAF1.2 is catalytically primed for signaling in the RRHCC complex, thereby reflecting a partial release of inhibitory constraints on RAF1 and potentially contributing further to its activation. These findings indicate that the stepwise regulation performed by the HSP90-CDC37 chaperone system could influence kinase or phosphatase activities by regulating kinase domain folding. Therefore, this maturation process may enable the phosphorylation/dephosphorylation events required to activate RAF1-driven MAPK signaling following RAF1 release from the chaperone system, subsequent assembly of its N-and C-lobes, and homo-or heterodimer formation through 14-3-3 adaptor proteins.

### Structure of the RHCp23 complex

While the structure of p23 in complex with HSP90 and a typical client, the glucocorticoid receptor (GR), has been determined^40^, the RHCp23 structure provides unprecedent mechanistic insight into how two functionally distinct cochaperones—CDC37, which mediates client recognition^41^, and p23, which modulates HSP90 ATPase activity ^42,43^—synergize to facilitate RAF1 folding (Fig. 3A-C, Fig. S2, SI3, Table S2, STAR Methods). p23 consists of a folded CHORD and Sgt1 (CS) domain and a largely unstructured tail region (Fig. 3A). The protein forms a complex with the closed client-bound state of HSP90 and has been shown to partially inhibit its ATPase activity^44^ ^43^. We identified two classes, one focused in the CDC37-MD (RHCp23_c) and another (RHCp23_p) displaying improved density for p23 (Fig. 3A-C).

**Figure 3.**
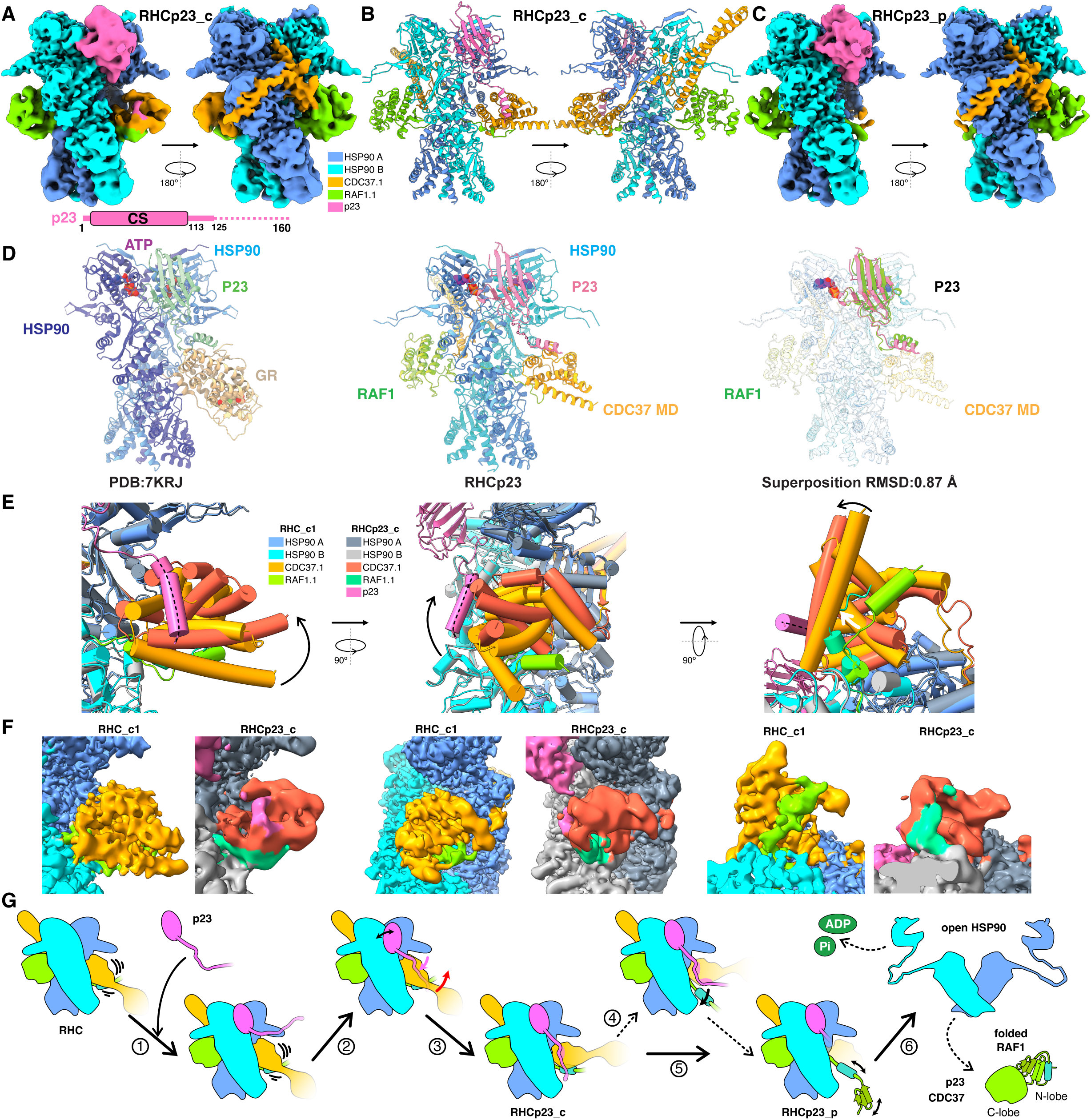
Cryo-EM structure of the RHCp23 complex. **(A)** Two views of the cryo-EM density map of class RHCp23_c with the domain organization of p23. **(B)** Ribbon representation of RHCp23_c in two orientations matching (a). **(C)** Cryo-EM map of class RHCp23_p. **(D)** Side-by-side view of the complexes of GR and RAF1 with HSP90-CDC37 showing the C-lobe of the RAF1 kinase domain in the opposite site of the GR complex while the MD of CDC37 is positioned in the same area. Detailed view of the superimposition of the complexes depicting the location of the P23 protein. **(E)** Comparison of CDC37-MD orientation in RHC_c1 and RHCp23_c. Ribbon diagrams of both structures in three perpendicular views; CDC37-MD movement is indicated by black arrows. The dashed line on top of the p23 helical section indicates the proposed modeling. **(F)** Corresponding cryo-EM maps of these regions, colored by protein. Additional density on CDC37-MD is shown in magenta. **(G)** Model for p23-assisted release of RAF1 from CDC37-MD in RHC. p23 first binds at the HSP90 dimer interface (1), enabling its flexible tail to contact CDC37-MD (2). This interaction tilts CDC37-MD (3), promoting release of the captured RAF1 N-lobe (5). Alternatively, the p23 tail may compete with the partially unfolded RAF1 N-lobe and block its binding (4). After ATP hydrolysis, the HSP90 dimer opens and releases RAF1 (6).

**Figure 4.**
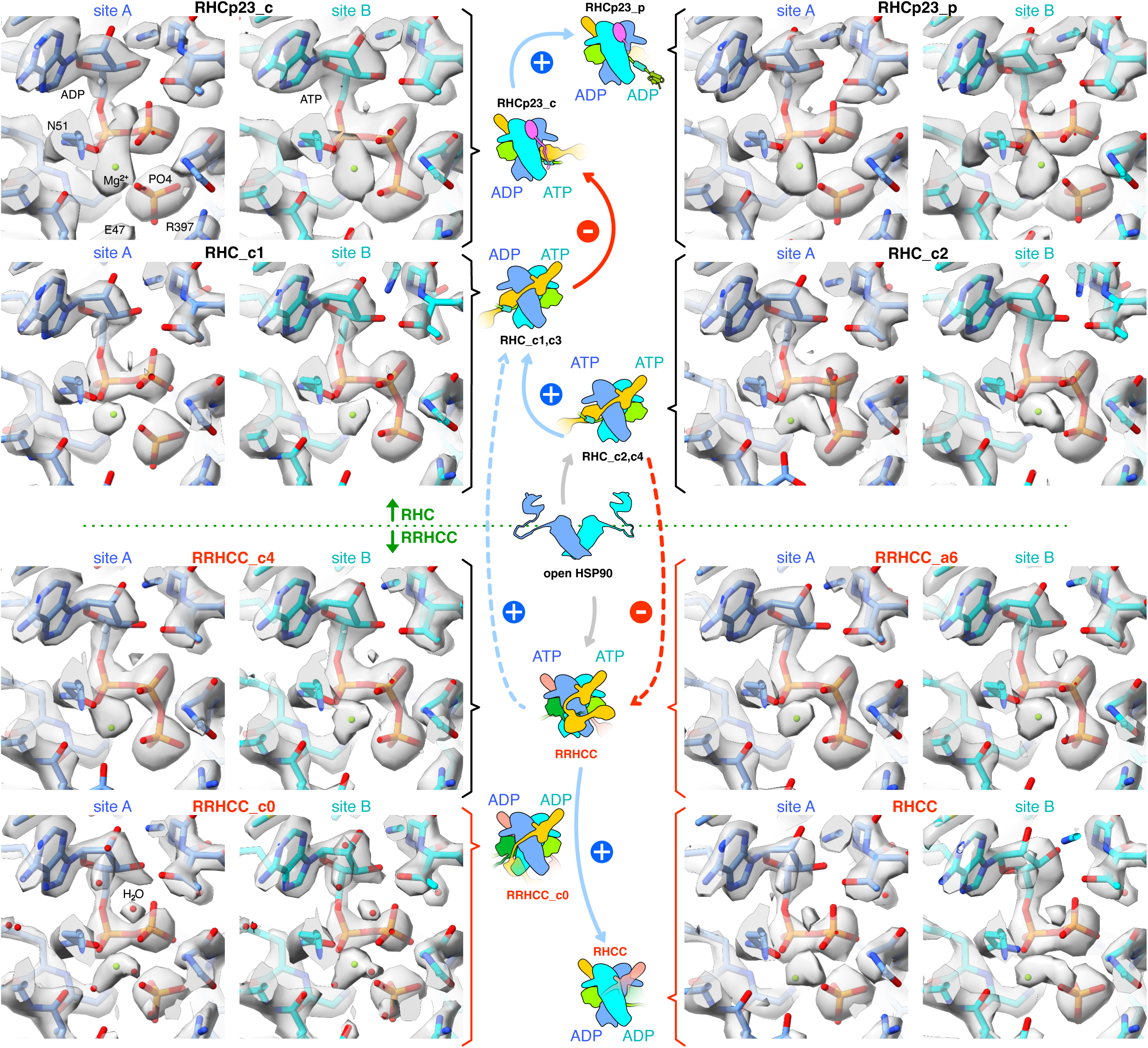
ATP hydrolysis and conformational changes in the RAF1-chaperone complexes. Composition of the HSP90 ATP-binding sites in the different cryo-EM maps (see also SI4). Each panel shows the density for a given site as a transparent surface, together with the modeled ligands at that site (phosphate is labeled as PO4). The central schematic summarizes the ATP-hydrolysis trajectory along the RHC and RRHCC assemblies, starting from an open HSP90 dimer. The upper branch corresponds to the RHC pathway, and the lower branch to the RRHCC classes. Blue arrows indicate increased HSP90 ATPase activity, red arrows indicate decreased activity, and dashed lines denote alternative transitions between the RHC and RRHCC pathways.

In the human GR-maturation complex (GR/HSP90/p23), the GR is threaded through the HSP90 lumen, with its ligand-binding domain folded and bound to the ligand, while p23 directly stabilizes native GR using a C-terminal helix^40^ in the overall disordered tail (Fig. 3D). Our structure shows that, unlike GR, p23 does not interact with RAF1, which is threaded through the clamp. In addition, extra density can be observed in the vicinity of CDC37-MD. Although this density is of low resolution, it is consistent with the presence of the flexible carboxy-terminal tail of p23 (Fig. 3E-F). The superposition of the RHCp23 complex with the GR/HSP90/p23 complex topologically supports this assignment in the neighborhood of CDC37-MD (Fig. 3D), suggesting that p23 employs its C-terminal tail as in the GR/HSP90/p23^42^ complex. As previously observed, the CDC37-MD is rather flexible and has been shown to bind an unfolded section of the kinase N-lobes of RAF1^24^, CDK4^33^ and the oncogenic BRAF^V600E^ mutant^28^ when assembled into complexes with HSP90. The dual interaction of p23 with HSP90 and the CDC37-MD could induce a tilt in CDC37-MD, disrupting its association with the unfolded N-lobe of RAF1 in an RHC complex (Fig. 3G). This conformational change would facilitate the release of the kinase. Furthermore, the possible interaction of the flexible tail with the CDC37-MD would prevent rebinding of the previously released unfolded N-lobe of RAF1 (Fig. 3G).

### The RHCp23 complex displays different nucleotide species

The changes induced in the structural elements of the NTD/MD interface of the HSP90 protomer, which are involved in ATP hydrolysis, are similar to those previously reported^40,42^. However, in contrast to the GR/HSP90/p23 structure, which displays ATP molecules in both ATPase domains, both γ-phosphates are cleaved in the RHCp23_p map, while only the ATP in site A is hydrolyzed in the RHCp23_c map (Fig. 4, and SI4 for comparison). p23 might delay ATP hydrolysis by HSP90 by inhibiting phosphodiester hydrolysis or by stabilizing the release of ADP+Pi. Our data suggest a two-step role for p23. First, p23 may stabilize the RHC complex and temporarily prevent hydrolysis of the second ATP (site B). Then, after the ATP in site B is finally hydrolyzed, p23 may help detach the αC helix from CDC37-MD, thereby accelerating HSP90 opening and promoting RAF1 release. Taken together, our findings show that p23 and CDC37 can coexist and operate in the same assembly during client maturation. This co-occupancy indicates tight coordination between client recruitment (CDC37) and ATPase regulation and substrate release (p23) (SI4B), suggesting that cochaperone pairing may be especially critical for complex clients like RAF1, which require fine-tuned regulation of domain unfolding/refolding.

### ATP binding and hydrolysis regulates RAF1-chaperone complexes

ATP binding triggers conformational changes in HSP90 that stabilize metastable client proteins like RAF1, helping them mature properly and avoiding misfolding or degradation^18^.

Our structural analysis across different RAF1-chaperone complexes and their ATP binding sites suggests that the cycle likely starts when HSP90 binds two ATP molecules and one RAF1–CDC37 complex to form an RHC complex (Fig. 4, SI4A). The RHC can either quickly recruit a second RAF1–CDC37 to form an RRHCC complex, thereby blocking ATP hydrolysis at both HSP90 sites, or proceed to hydrolysis at site A (RHC_c1, c3). This prevents RRHCC assembly and facilitates the assembly of p23. When only one RAF1–CDC37 is bound, site A can hydrolyze ATP, after which the RHC complex (RHC_b) becomes competent to bind p23, delaying hydrolysis. The resulting RHCp23_c states share the same nucleotide pattern and protect site B from hydrolysis (RHCp23_c). Later, when both sites contain ADP+Pi (RHCp23_p), the RAF1 N-lobe begins to be released for folding (Fig. 3G). In the case of the RRHCC complex, the hydrolysis of both ATPs leads to a state (RRHCC_c0) in which the N-lobe of RAF1.2 has been released and an RHCC complex forms as a final stage towards complex disassembly. Alternatively, RRHCC complexes could convert back to an RHC-like state through selective hydrolysis at site A (RHC_c1, c3), coupled to release of RAF1.2 and CDC37.2 (Fig. 4A, SI4). Overall, the number of RAF1–CDC37 copies bound to HSP90 and the precise timing of ATP hydrolysis at the two sites would function as checkpoints that control whether RAF1 is stabilized, allowed to fold, or released for downstream signaling (SI4B). Collectively, these observations support a model in which ATP binding and hydrolysis by HSP90 tightly coordinate the assembly, remodeling, and disassembly of RAF1 chaperone complexes.

### RAF1 and BRAF^V600E^ form homodimers when in complex with HSP90-CDC37

The interplay between RAF family dimerization mediated by 14-3-3 proteins and the stabilization of RAF1 and other RAF isoforms clients of the HSP90-CDC37 chaperone system is crucial for the regulation of RAF activity, as the final formation of RAF dimers with 14-3-3 is essential for kinase activation and efficient MEK phosphorylation ^13,45–47^.

Therefore, understanding how RAF dimers assemble and stabilize remains an important mechanistic question. The identification of the RRHCC complex prompted us to investigate whether other RAF family members, which are known clients of HSP90-CDC37, can form this homodimeric intermediate assembly observed with RAF1.

To address this question, we conducted pull-down experiments after transient expression of HaloTag-KRAS^G12V^, CDC37 and different RAF isoforms under the same conditions used to obtain the RRHCC complex. The different RAF isoforms combined the strep and V5 tags in Expi293F cells (Fig. 5A, STAR Methods). Our panel included well-characterized RAF clients of the HSP90-CDC37 system, such as BRAF^V600E^ and ARAF, along with the kinase-dead oncogenic BRAF^D594A^ and wild-type BRAF, which seems to interact weakly with HSP90-CDC37^10,48^. The different tag combinations enabled both the purification of these assemblies and the detection of the components of the chaperone-associated complexes via western blotting with tag-specific antibodies (Fig. 5A-B, Fig. S9A, STAR Methods). In addition, following pull-down, we identified the presence of individual proteins using mass spectrometry and performed a relative quantification of each assembly by comparing the abundance of unique peptides corresponding to each tagged protein within the complex (Fig. 5B, STAR Methods). To provide a comparison with other kinase clients beyond the RAF family, we included CDK4, a well-established interactor of the HSP90-CDC37 system that does not require dimerization for activation^33^, as a negative control.

**Figure 5.**
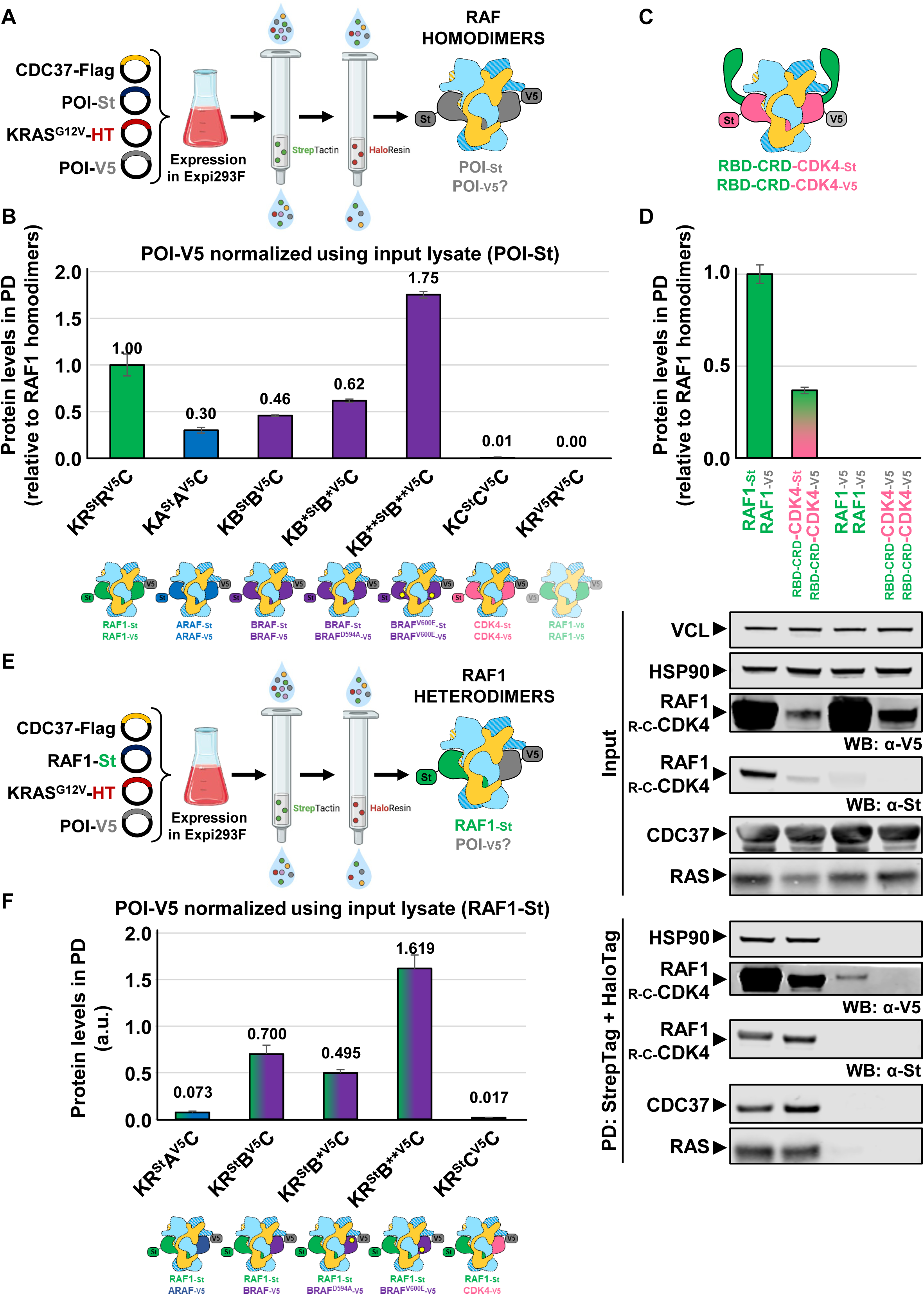
Dimerization of RAF family proteins in the context of the RRHCC complex. **(A)** Schematic representation of the experimental workflow used to detect RAF homodimers by sequential StrepTag and HaloTag purifications from clarified cell lysates. **(B)** Quantification of dimer formation by mass spectrometry, based on unique V5-tagged peptides of the protein of interest (POI) detected in the eluates. Graphs show mean protein levels normalized to POI-Strep peptides and are expressed relative to RAF1 homodimer levels. The plot represents the results from two independent experiments, and error bars indicate standard deviation. **C)** Illustration of the RBD–CRD–CDK4 chimeric protein forming a 2:2:2 stoichiometric complex with HSP90 and CDC37. **D)** *Top*. Quantification of dimer formation as described in (C). *Bottom*. Western blot analysis of input lysates and pull-down samples from the experiment. Solid arrowheads indicate the migration positions of the relevant proteins. **(E)** Schematic representation of the experimental workflow used to detect RAF1 heterodimers by sequential StrepTag and HaloTag purifications from clarified cell lysates. **(F)** Quantification of dimer formation by mass spectrometry, based on unique V5-tagged peptides of the protein of interest (POI) detected in the eluates. Graphs show mean protein levels normalized to RAF1-Strep peptides present in the input lysate and are expressed as arbitrary units. The plot represents the results from two independent experiments, and error bars indicate standard deviation. The schematic representations below panels **B)** and **F)** depict the composition of the analyzed complexes: HSP90 (light blue), CDC37 (yellow), RAF1 (green), ARAF (dark blue), BRAF (purple), CDK4 (pink), and KRAS (not shown). Abbreviations: St, StrepTagII; HT, HaloTag; POI, protein of interest. Complex notations: KR^St^R^V5^C=HaloTag-KRAS^G12V^ + RAF1–StrepTagII + RAF1–V5 + CDC37–Flag; KX^St^X^V5^C = HaloTag-KRAS^G12V^ + X–StrepTagII + X–V5 + CDC37–Flag, where X represents: A = ARAF, B = BRAF, B* = BRAF^D594A^, B** = BRAF^V600E^, C = CDK4. See also Figure S9.

Our results demonstrate that the formation of homodimeric assemblies within the HSP90-CDC37 complex is selective and varies significantly across RAF isoforms. Notably, BRAF^V600E^ was the most enriched dimeric species in our analysis (175% relative to RAF1) (Fig. 5B), in agreement with emerging evidence indicating that this oncogenic mutant can adopt a dimeric state in the context of the HSP90-CDC37 complex^28,49^. The kinase-dead BRAF^D594A^ mutant also formed substantial homodimers (62%), whereas wild-type BRAF and ARAF displayed more modest associations at 46% and 30%, showing that either they interact weakly with HSP90^28,48^ (BRAF) or do not have an efficient formation of the homodimeric assembly with the chaperone system (ARAF). By contrast, CDK4, a canonical client of HSP90-CDC37 that does not require dimerization, was present at only 1%, highlighting the specificity of this chaperone-specific dimer formation among RAF clients (Fig. 5B). These data reinforce a model in which the dimerization of RAF isoforms within the HSP90-CDC37 complex is a selective process and may represent a critical intermediate for the stabilization and functional maturation of certain RAF family members.

### The RRHCC complex is exclusive to RAF kinases

Given that CDK4 does not interact with KRAS and the Strep-Halo affinity purification will select associations involving the GTPase, the absence of CDK4-dimeric species is not unexpected. This led us to investigate whether the assembly of such chaperone-associated dimeric intermediates is a distinctive regulatory feature of RAF family members. A plausible explanation for this specificity could be the presence of the CR1, which includes the RBD and the CRD, unique to RAF proteins and absent in CDK4 and other kinase clients of the HSP90-CDC37 system. To experimentally test this hypothesis, we engineered a chimeric protein substituting the kinase domain of RAF1 by CDK4 generating an RBD-CRD-CDK4 fusion protein (Fig. 5C, STAR Methods). This fusion protein was co-expressed with HaloTag-KRAS^G12V^, as we did with RAF family members, to assess whether the presence of RAF-specific regulatory elements could promote the formation of similar dimeric intermediates within the chaperone complex.

The analysis showed that the presence of the RBD and CRD domains induced the formation of dimers associated with the HSP90-CDC37 complex (Fig. 5D). To assess whether this effect depends on the mere presence of KRAS^G12V^ or requires a direct interaction of KRAS^G12V^ with the kinase, we performed a control experiment in which CDK4 was transfected with two different tags (His and V5), along with HaloTag-KRAS^G12V^ and CDC37-Strep. However, the HaloTag of KRAS was not used to pull down proteins; instead, CDC37 was first pulled down, followed by His-tagged CDK4 to assess complexes where the chaperones are involved (Fig. S9B-C). We did not detect CDK4 homodimers in this setup, supporting the idea that interaction with KRAS^G12V^ facilitates the formation of the higher-order complex consisting of two molecules each of HSP90, CDC37, and the associated kinase. The influence of RAS in inducing RAF dimerization with 14-3-3 has been previously observed^47^. Our experiments also suggest that RAS-GTP^6,8^ affects RAF dimerization by increasing the local concentration of the different kinase isoforms associated with HSP90-CDC37. Quantification of the chaperone-bound complexes revealed that the RBD-CRD-CDK4 chimera formed homodimers at a substantial level, ∼40% relative to RAF1 homodimers, indicating that the presence of the RBD-CRD module is sufficient to promote dimerization within the HSP90–CDC37 complex. Notably, all tested RAF family members, including both wild-type and oncogenic mutants, also assembled into homodimeric complexes to varying extents, ranging from 30% (ARAF) to 175% (BRAF^V600E^).

Hence, these findings suggest that while the RBD-CRD domains facilitate dimerization, the full extent of complex formation is likely influenced by additional features, including the intrinsic folding stability of the kinase domain and its affinity for the chaperone system. We propose that the association of KRAS^G12V^ with the RBD, in combination with a thermodynamically wobbly RAF1 kinase domain, triggers the formation of the intermediate complex containing a 2:2:2 stoichiometry with HSP90-CDC37 to regulate kinase maturation. Collectively, these findings show that the HSP90-CDC37 chaperone system articulates a singular maturation mechanism for the high-affinity clients of the RAF family. This stabilization pathway differs from that of classical kinase clients, such as CDK4, which do not incorporate multiple protomers into the assembly.

### HSP90-CDC37-associated RAF1 forms heterodimers with other RAF isoforms

Heterodimerization of RAF kinases is a central mechanism regulating the MAPK pathway in both normal and oncogenic contexts^1^ ^13^. The 14-3-3 proteins facilitate and stabilize these dimers by binding phosphorylated serine residues and ensuring proper conformational states for activation^50^. While RAF1 is thought to be the dominant RAF activated by oncogenic KRAS, its kinase activity is not required for KRAS-driven tumor formation^14,51^. However, its ability to dimerize is essential^51^, and heterodimer formation often enhances signaling efficiency^52^.

Having demonstrated that RAF1 and BRAF^V600E^ can each form homodimeric assemblies within the HSP90-CDC37 complex, we next explored whether the different RAF isoforms similarly associate to build heterodimers with RAF1 when stabilized by the HSP90-CDC37 chaperone system (Fig. 5E-F, Fig. S9D, STAR Methods). Remarkably, we detected heterodimeric assemblies of RAF1 mainly with BRAF^V600E^ and to a lesser extent with BRAF and BRAF^D594A^ (Fig. 5F). By contrast, heterodimers with ARAF showed minimal abundance in agreement with the lower observed homodimerization (Fig. 5B). When the assay was conducted to evaluate the presence of mixed complexes including RAF1 and other kinase clients, i.e. CDK4, we were unable to detect the heterodimeric association, indicating again that the formation of these assemblies is restricted to certain RAF isoforms (Fig. 5F, Fig. S9D). This model might be especially critical for the maturation of RAF1 and BRAF^V600E^, which strongly depend on chaperone assistance for their stability and functional activation^24,28^. In both cases, we can identify intermediate homo-and heterodimeric kinase-chaperone-cochaperone-client complexes.

### RAF homo-and heterodimers modulate MEK phosphorylation and cellular proliferation

Taken together, our findings indicate that the HSP90–CDC37 chaperone system not only stabilizes RAF kinases but also actively primes them for functional activation by promoting a stepwise folding process that ends in dimerization-competent conformations with 14-3-3 poised for subsequent signaling through the MAPK pathway^12,13,15,24,31^. This led us to explore whether RAF molecules emerging from this chaperone-assisted assembly would exhibit differential signaling capacities depending on their dimeric partners. To investigate the ability of individual RAF isoforms (homodimers) and their pairwise combinations with RAF1 (heterodimers) to reconstitute MAPK signaling (Fig. 6A), we utilized *ARaf^Y/lox^*; *BRaf^lox/lox^*; *Raf1^lox/lox^* mouse embryonic fibroblasts (MEFs), in which ablation of RAF family members can be achieved upon adenoCRE infection (RAFless)^24^. Consistent with their biochemical assembly within the HSP90-CDC37 complex, ectopic expression of RAF1 and BRAF^V600E^ led to the strongest activation of pMEK. BRAF and BRAF^D594A^, which displayed lower homodimerization efficiency, showed weak levels of MEK activation (Fig. 6A). Co-expression of RAF1 with BRAF^V600E^ or with the kinase-impaired BRAF^D594A^ mutant led to robust MEK phosphorylation (Fig. 6A). This is particularly notable in the case of BRAF^D594A^, suggesting a paradoxical transactivation mechanism^3,53^ in which RAF1 serves as the active kinase partner in a heterodimer, compensating for the catalytic inactivity of BRAF^D594A^. Similarly, RAF1 co-expression with wild-type BRAF also resulted in MEK activation, likely due to enhanced heterodimerization and chaperone-assisted complex stabilization.

**Figure 6.**
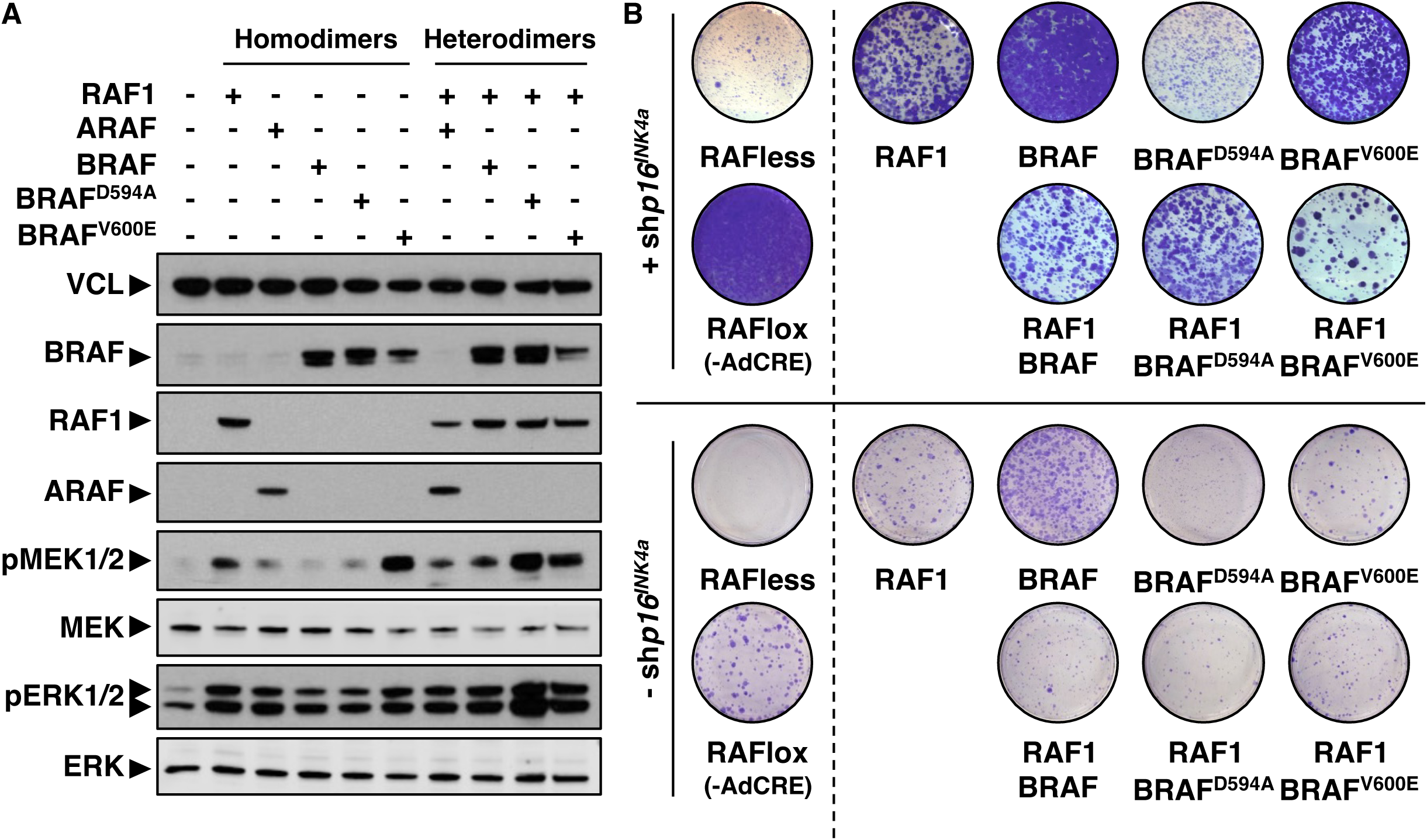
Ectopic expression of RAF dimers differentially restores MAPK signaling and proliferation in RAFless MEFs. **A)** Western blot analysis showing activation of the MAPK pathway in RAF-deficient MEFs. RAFlox MEFs were transduced in parallel with retroviruses expressing the indicated RAF isoforms (ARAF, BRAF, BRAF^D594A^, or BRAF^V600E^), either alone (homodimers) or in combination with lentiviruses expressing RAF1 (heterodimers). Subsequently, cells were infected with adenoviruses encoding CRE recombinase to induce the deletion of all endogenous RAF genes. Phosphorylated MEK1/2 (pMEK1/2) levels were assessed by immunoblotting and quantified relative to parental RAFless cells, after normalization to the loading control. **B)** Colony formation assay of RAFless MEFs expressing the indicated RAF isoforms, with or without a shRNA targeting p16^INK4a^ to prevent cellular senescence. Ablation of the endogenous RAF genes was achieved as described in panel (a). Representative crystal violet–stained plates from one experiment are shown.

Next, we tested whether these complexes enable proliferation in the absence of endogenous RAF proteins. We performed a colony-forming assay using RAFless MEFs ectopically expressing the combinations of the RAF isoforms observed to assemble the 2:2:2 intermediate (Fig. 5B, 5F, Fig. S10, STAR Methods). To avoid cell senescence-like growth arrest via *p16^INK4a^*upregulation^54,55^, we co-infected RAFless MEFs with a shRNA targeting *p16^INK4a^* (STAR Methods). In this cellular context, both RAF1 and BRAF^V600E^ restored proliferation. Remarkably, the expression of BRAF^D594A^ did not elicit proliferation, whereas the RAF1-BRAF^D594A^ combination became competent to support growth, likely due to paradoxical transactivation of RAF1^3,53^ (Fig. 6B). These findings suggest that the assembly of homo-and heterodimers within the HSP90–CDC37 leading to kinase activation influences the signaling output and thereby cell proliferative competence.

### An integrated model for RAF maturation

Our data support a comprehensive RAF1 maturation cycle mediated by the HSP90-CDC37 chaperone machinery, which is extendable to other HSP90-CDC37-dependent RAF isoforms. RAF1-CDC37 associates with HSP90 through multiple routes requiring ATP binding (Fig. 4, 7A). Single RAF1-CDC37 complexes form RAF1-HSP90-CDC37 (RHC) assemblies (step 1a), while two RAF1-CDC37 complexes associate simultaneously or sequentially to assemble RAF1-RAF1-HSP90-CDC37-CDC37 (RRHCC) complexes (steps 1b-1c). In RHC assemblies lacking a second RAF1-CDC37 complex, p23 binds HSP90, regulating nucleotide hydrolysis and promoting RAF1 αC helix release (steps 2a-3a). Successful N-lobe formation upon HSP90 opening yields fully folded kinase domains (step 4a). In RRHCC complexes, ATP hydrolysis drives RAF1.2 αC helix release from CDC37.1-MD (step 2b), producing RHCC intermediates (step 3b) that either undergo complete disassembly (step 4b) or, following CDC37.2 dissociation, bind p23 and proceed through RHC-like pathways (step 4c). Several steps of this maturation occur at the membrane context where RAF1 engages KRAS through its RAS-binding domain (RBD) (Fig. 7B). The initial composition of membrane-localized KRAS-RRHCC-14-3-3 complexes could predetermine the formation of homo-or heterodimeric active RAF complexes (RAF1-RAF1, RAF1-BRAF, or RAF1-ARAF). The interaction of 14-3-3 with phosphorylated sites S259 and S621 locks the kinases in complex with HSP90 and directs them toward active homo/heterodimer formation. Simultaneously, RAF1-KRAS interaction anchors all complexes to the membrane, maintaining local concentration. The joint action of KRAS and 14-3-3 on chaperone-bound RAF1 may enhance proper kinase folding by promoting N-lobe dimerization, creating a coordinated transition from chaperone-dependent maturation to 14-3-3-stabilized active dimers competent for MEK phosphorylation.

**Figure 7.**
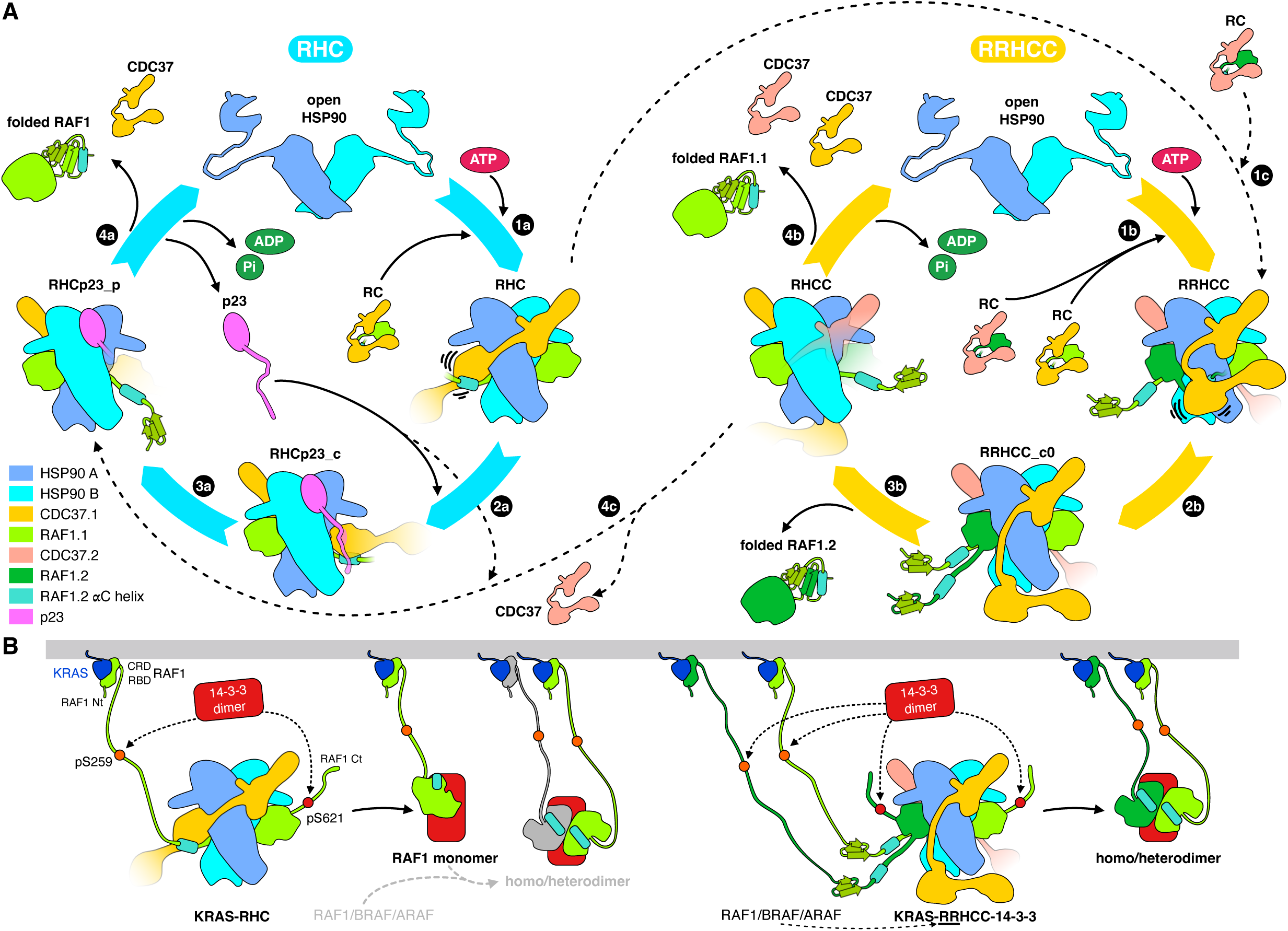
Model of RAF1 maturation by the HSP90-CDC37 system. **A)** Scheme of the RAF1 stepwise folding mechanism mediated by HSP90-CDC37. **B)** Assembly of the homo-and heterodimers for RAF1 activation at the membrane context. For clarity, the autoinhibited RAF1 monomer and MEK substrate are not depicted (see main text for details).

## DISCUSSION

The results presented here reposition the HSP90–CDC37 system from a passive stabilizer of RAF clients to an active organizer of RAF signaling competence. By resolving multiple RAF1-containing assemblies, we capture snapshots of a maturation cycle that couples folding, nucleotide-state control, post-translational remodeling, and dimer readiness into a single, integrated mechanism (Fig. S2, Fig.7A-B). A central conclusion is that RAF1 maturation proceeds through an *asymmetric, stepwise* pathway that is qualitatively different from the canonical “single-client” logic described for many HSP90–CDC37 kinase clients^56,28,33^.

The RRHCC architecture suggests that HSP90-CDC37 does not simply hold RAF1 in a uniform unfolded conformation but partitions the folding problem and imposes order on the αC-specific regulatory element while delaying completion of the ATP-binding architecture (Fig. 4). In doing so, HSP90–CDC37 appears to implement a *conformational checkpoint* that prevents premature adoption of an active kinase-competent state keeping the “histidine anchor” region secluded until the appropriate assembly context is achieved.

Mechanistically, our work identifies a mold-like interface created by CDC37 and the HSP90 MD that stabilizes the partially folded αC helix and registers it via the conserved “histidine anchor” (Fig. 1D-L). The functional impact of this node is emphasized by the mutational analysis, which showed that disrupting anchor contacts markedly depletes RAF1 from isolated complexes (Fig. 2E). This is consistent with destabilization and degradation of improperly matured kinase. Conversely, modifying the acidic surface at CDC37-MD opposes helix extension and increases RAF1 recovery (Fig. 2F). This result indicates that this region acts as a built-in brake—stabilizing an αC “seed” while preventing full helix propagation until the system commits to release. Taken together, these observations suggest that HSP90–CDC37 can influence kinase activity by selectively stabilizing a regulatory element that is both structurally fragile and functionally decisive, thereby shaping not only folding success but also the downstream allosteric landscape that governs activation. In parallel, emerging therapeutic strategies seek to disrupt oncogenic kinase signaling by destabilizing HSP90–CDC37 interactions using small-molecule inhibitors, peptides, peptidomimetics, and natural products. ^57^. In this context, our findings offer a framework for selectively targeting RAF isoforms.

The hypothesis that cochaperone pairing provides temporal control over this checkpoint is supported by the RHCp23 structure (Fig. 3A-C), which demonstrates that p23 can coexist with CDC37 on the same closed HSP90 clamp. Binding of p23 likely biases the ATPase cycle to prolong a client-bound state while promoting productive release when hydrolysis progresses. This arrangement suggests a coordinated action in which CDC37 confers kinase selectivity and supports the “molded” intermediate, whereas p23 tunes dwell time and opening propensity, limiting futile cycles of rebinding. Collectively, the data suggest that cochaperones could be viewed as modules that adjust the energetic and kinetic gates of HSP90^58^ to match the maturation requirements, especially for complex clients like RAFs.

The nucleotide occupancy across structural classes indicates that ATP binding and asymmetric hydrolysis act as decision points that route complexes toward alternative outcomes. Differences in hydrolysis state correlate with CDC37-MD variability and with transitions between single-client and double-client assemblies, implying that client stoichiometry itself can modulate ATPase timing. This provides a plausible mechanism by which HSP90 can “sense” the assembly state of RAF1 and coordinate progression from stabilization to release (Fig. 4, 7A). In this view, ATP turnover is not merely the engine of chaperone cycling; it is a programmable gate that couples chemical state to client maturation status.

Finally, phosphoproteomics indicates that within the RRHCC context, RAF1 is selectively enriched for activating phosphorylations while inhibitory sites are reduced (Fig. S1H). These results suggest that the chaperone environment biases the phosphorylation landscape by regulating kinase accessibility to modifying enzymes and facilitating *pre-dimerization*, which primes RAF to form active homo-and heterodimeric kinase assemblies modulating signaling (Fig. 5, 6). Importantly, the ability of a RAF regulatory module (RBD–CRD) to confer dimeric behavior on an otherwise “canonical” client underscores that this pathway is not simply an intrinsic property of the kinase domain; it emerges from the intersection of RAF-specific regulatory architecture, RAS-driven concentration effects, and chaperone-mediated folding control.

Overall, these findings establish the HSP90-CDC37 chaperone system as an active regulator rather than passive stabilizer of RAF kinase maturation. These processes collectively determine signaling competence. This mechanistic framework extends beyond RAF1 to other RAF family members and highlights potential therapeutic vulnerabilities. Disrupting RAF-chaperone interfaces or blocking predimerization may selectively inhibit aberrant RAS-RAF signaling without compromising essential kinase functions dependent of HSP90-CDC37. More broadly, the coupling of chaperone-mediated folding to signaling competence represents a paradigm shift in understanding how protein quality control systems dictate signal transduction logic, with implications extending well beyond the MAPK pathway.

## LIMITATIONS OF THE STUDY

The RAF-less MEF assays provide functional context for chaperone-associated RAF assemblies but do not establish specific chaperone-bound intermediates as the unique drivers of growth rescue. Although homo-and heterodimer assembly within the HSP90–CDC37 system is supported by pull-down assays, concurrent formation of non-client RAF isoforms facilitating proliferation cannot be excluded (e.g for BRAF). In addition, structural elements implicated in chaperone-assisted maturation also contribute to kinase activation, making it difficult to disentangle effects on folding from effects on signaling output. Future approaches capable of selectively perturbing individual maturation or dimerization states will be needed to resolve these relationships.

## ACKNOWLEDGMENTS

We thank the Danish cryo-EM National Facility at the University of Copenhagen for support during data collection supported by grant NNF14CC0001, NNF24SA0098829 to the Novo Nordisk Foundation Center for Protein Research (CPR). We also thank Dr. J. Boskovic and of the EM unit at CNIO for negative stain analysis of the sample. J.R.C-T is supported by an MSCA-HE postdoctoral fellowship (#101065213). This project was supported by La Caixa Foundation (ID 100010434) with fellowship codes LCF/BQ/DR22/11950011 and LCF/BQ/DR23/12000028. L.d.l.P.-O. is supported by a predoctoral fellowship from the Spanish Ministry of Universities (FPU21/04678). G.M. and J.V.O. are part of CPR, which is supported financially by the Novo Nordisk Foundation (NNF14CC0001, NNF24SA0098829). This work was also supported by the ERC-AdG 101096548 (INTETOOLS), NNF0024386, NNF17SA0030214, and NNF18OC0055061 grants to G.M, who is a member of the Integrative Structural Biology Cluster (ISBUC) at the University of Copenhagen. The authors acknowledge the Doctoral School of Molecular Biosciences of the Faculty of Sciences at Universidad Autónoma de Madrid for academic support.

## AUTHOR CONTRIBUTIONS

This study was conceptualized by GM and S.G-A. G.A. P.M., J.R.C-T., M.B., S.G-A., and G.M. designed biochemical experiments. G.A., J.R.C-T. and S.G-A. set up the purification protocol. G.A., L.d.l.P.O., and L.L.R created the mutants and performed biochemical experiments. G.A., L.d.l.P.O. and C.G.L. performed tissue culture experiments. E.Z., M.I., J.R.C-T., L.v.d.H., and J.V.O. performed mass spectrometry and analysed the results. G.A., J.R.C-T. and P.M. performed the final purification, prepared cryo-EM grids and collected the cryo-EM images. P.M. performed cryo-EM data processing and built the structural models.

P.M. and G.M. proceeded with cryo-EM map and structure analysis. The global results were discussed and evaluated by all authors. G.M. and S.G-A. coordinated and supervised the project and wrote the manuscript with input from all the authors.

## COMPETING INTEREST

GM is a stockholder of Ensoma and has been consultant for Orbis Medicines. MB declares that is co-founder and shareholder of Vega Oncotargets S.L. The rest of the authors declare no conflict of interest.

## STAR METHODS

Detailed methods include the following:

- RESOURCE AVAILABILITY

.- Lead contact
.- Material availability
.- Data and code availability
- EXPERIMENTAL MODEL AND SUBJECT DETAILS

.- Cell lines
- METHOD DETAILS

.- *Plasmid cloning, expression and sample preparation*
.- *Cryo-EM Sample preparation and data collection*
.- *Cryo-EM data processing and conformational analysis*
.- *Model building and refinement*
.- *Phosphoproteomics*
.- *Mass spectrometry quantification*
.- *Site-directed mutagenesis and pulldowns from RAF1 and CDC37 mutants*
.- *Pulldowns from CDC37-Strep and his-tagged kinases*
.- *Western blot analysis and antibodies*
.- *Heterodimer and homodimer formation assay*
.- *Phosphorylation in RAFless MEFs*
.- *RAFless proliferation assay*

## SUPPLEMENTAL INFORMATION

Figs. S1 to S11

Supporting Information 1-4

Tables S1 Phosphorylation of RAF1-chaperone complexes Table S2 CryoEM data collection and refinement

Table S3 Oligonucleotides, cell lines, etc.

## STAR METHODS

### RESOURCE AVAILABILITY

#### Lead contact

Further information and requests for resources and reagents should be directed to and will be fulfilled by the lead contact.

#### Materials availability

Reagents generated in this study are available from the lead contact upon request with completed material transfer agreements.

#### Data and code availability

- Protein Data Bank (PDB) and Electron Microscopy Data Bank (EMDB) identification numbers for the cryo-EM structures and maps reported in this manuscript are available as of the date of publication. Identification numbers are listed in the Table S2. Phosphoproteomic data have been deposited in the PRIDE database with project accession code PXD065622 (See Table S1).
- Any additional information required to reanalyze the data reported in this paper is available from the lead contact upon request

#### Experimental Model and Subject Details

##### Cell lines

“Raflox” Mouse Embryonic Fibroblast (MEF) cell lines were generated at Barbacid Laboratory. Cells were cultured in DMEM (Gibco) supplemented with 10% FBS (Gibco) at 37 □C and 5% CO_2_ in a humidified incubator. Cells were suspended using Trypsin-EDTA solution (Gibco) and split every 2 to 3 days.

## Method Details

### Plasmid cloning, expression and sample preparation

The complete coding sequence for human KRAS (Accession No. AF493917.1) containing the activating G12V mutation and fused to an HA tag was cloned into the mammalian expression vector pFN21A HaloTag® CMV Flexi® Vector (Promega, Madison, WI, USA) following the manufacturer’s instructions. The resulting plasmid, pFN21A-Halo-KRAS^G12V^, includes the HaloTag sequence (Accession No. EU621374) at the N-terminus of KRAS^G12V^. Additional plasmids, including pcDNA3-RAF1-Strep and pCMV-CDC37-Myc-DDK, were prepared as previously described^24^. All plasmids were sequence-verified. Expi293F cells (Thermo Fisher Scientific) were maintained and expanded according to the manufacturer’s protocol. Briefly, cells were cultured in Expi293™ Expression Medium (Thermo Fisher Scientific) at 37°C, 5% COe, and 120 rpm in PETG Erlenmeyer flasks (Corning) equipped with 0.2 μm vented caps and subcultured approximately every 3–4 days. For transfection, cells were diluted to a final density of 3×10e cells/mL in a total volume of 240 mL in 1 L flasks and transfected using the ExpiFectamine™ 293 Transfection Kit (Thermo Fisher Scientific) following the manufacturer’s protocol. A total of 240 μg of plasmid DNA (96 μg pcDNA3-RAF1-Strep, 96 μg pFN21A-Halo-KRAS^G12V^, and 48 μg pCMV-CDC37-Myc-DDK) was diluted in Opti-MEM™ I GlutaMAX™ (Thermo Fisher Scientific). Enhancers 1 and 2 (Thermo Fisher Scientific) were added 20 hours post-transfection. Cells were then incubated under standard conditions for an additional 48 hours before harvesting.

Two batches of transfected Expi293F cells (480 mL total culture volume) were harvested and resuspended in 40 mL of lysis buffer (20 mM Tris-HCl pH 7.5, 150 mM NaCl, 10 mM MgCle, 20 mM NaeMoOe, 0.1% Triton X-100) supplemented with cOmplete™ Mini EDTA-free Protease Inhibitor Cocktail (Merck), Phosphatase Inhibitor Cocktails 2 and 3 (Merck), and 1,000 U of benzonase nuclease. Lysates were incubated on ice for 10 minutes, followed by ultrasonic disruption (two pulses of 30 seconds at 20% amplitude). Lysates were clarified by centrifugation (13,000 rpm, 20 minutes, 4°C). The soluble fraction was subjected to two consecutive purification steps: (i) Strep-affinity chromatography (Strep-AC) and (ii) Halo-affinity chromatography (Halo-AC). For Strep-AC, 8 mL of Strep-Tactin® XT 4Flow High-Capacity resin (IBA Lifesciences) was packed into two 10 mL Pierce™ centrifuge columns (Thermo Fisher Scientific) and equilibrated with 12 column volumes (CV) of KRC buffer (20 mM Tris-HCl pH 7.5, 150 mM NaCl, 10 mM MgCle, 20 mM NaeMoOe). The soluble fraction was incubated with the resin on ice for 10 minutes. After washing with KRC buffer, KRC buffer supplemented with 5 mM biotin (IBA Lifesciences) was added, and the resin was incubated on ice for 5 minutes prior to elution. Elution with KRC buffer was performed twice. Eluted fractions were directly loaded onto 8 mL of HaloLink™ Resin (Promega) pre-equilibrated with KRC buffer containing 0.005% IGEPAL® CA-630. Samples were incubated for 1 hour on ice. After washing with the same buffer, proteins were eluted by digestion with Halo-TEV Protease (Promega) at a final concentration of 0.15 U/mL for 90 minutes on ice. The final eluate was concentrated using Amicon® Ultra-4 centrifugal filter units (10 kDa cutoff, Merck) to a protein concentration of approximately 1.5 mg/mL. One fifth of the concentrated sample was directly used for cryo-EM grid preparation. The remaining portion was loaded onto a Superdex® 200 Increase 3.2/300 gel filtration column (Cytiva) pre-equilibrated with KRC buffer, using an ÄKTA™ FPLC system (Cytiva Life Sciences). The pre-and post-gel filtration samples were analyzed by mass spectrometry.

### Cryo-EM Sample preparation and data collection

KRHC purified samples in KRC buffer (3 μl) were applied to grids (dataset A: UltraAuFoil R1.2/1.3 Au 300 mesh, dataset B: QuantiFoil R1.2/1.3 Cu 300 mesh, from QuantifoilMicroTools GmbH) previously glow-discharged in a Leica EM ACE200 for 60 s at 10 mA. Grids were plunge-frozen in liquid ethane pre-cooled with liquid nitrogen using a Vitrobot Mark IV (Thermo Fisher Scientific) with a blotting time of 4 s, 100% humidity, 4°C. The best grids were imaged at the Core Facility for Integrated Microscopy (CFIM – University of Copenhagen) on a Titan Krios G2 (300 kV, Thermo Fisher Scientific) equipped with a Falcon 4i direct electron detector and a Selectris X energy filter. Zero-loss images were collected using an energy filter slit width of 10 eV. Movies (Extended data Figure 3, Supp. Table 2) were collected using EPU v3.10.0.8733 at 165000x magnification (dataset A: 0.725 Å/pixel, 50.43 electrons/Å^2^ and 2.72 s exposure time; dataset B: 0.728 Å/pixel, 60 electrons/Å^2^ and 2.87 s exposure time), with nominal defocus from-0.8 to-2.0 μm in 0.2 μm intervals. As we previously identified a problem with the distribution of orientations of the RHC particle^24^, we collected approximately half of the data with a 30° tilting of the grids.

### Cryo-EM data processing and conformational analysis

Data from multiple samples of RAF1-chaperone complexes were collected and preliminarily analyzed. Two of them were selected because of the presence of special complexes, RRHCC and RHCp23 (dataset A and B respectively, Fig. S2). CryoSPARC v4.7.0 was used for image processing ^59^. Movie frames were motion-corrected and dose-weighted using patch motion correction, and CTF parameters were calculated by the patch CTF estimation job. Micrographs were initially picked using the blob picker job with the parameters of the previously described RHC complex^24^. Particles were extracted with a box size of 448 pixels (or downsampled to 224 pixels). We used initial 2D classifications to obtain a subset of particles for *ab initio* reconstruction. Although this initial map looked like the typical RHC particle, we were able to see differences in some of the 2D classes (Fig. S2B-D), which were further clarified with additional processing and classification. To solve the different levels of heterogeneity in the population, we used different classification methods in parallel: some unfocused (heterogeneous refinement and 3D classification) and other focused in specific regions (3D classification and 3D variability analysis, mainly CDC37-MD or CTD, and p23). Significantly, the different described ATP-hydrolysis states (Fig. 4, S3B, S4E, S5C, SI4) were indirectly isolated when trying to solve the CDC37/p23 heterogeneity and not the specific HSP90-NTDs. Consequently, the classifications highlighted a compositional and conformational interconnection of the complexes with the contents of the ATP binding sites of their HSP90 subunits. The contents of the ATP sites were assigned by comparison among all the structures of the coulombic densities at the catalytic sites while keeping uniform the values of the rest of HSP90 nearby regions (Fig. 4, SI 4). In this respect, RHC_b showed a particularly strong signal for Pi at site A, which we interpreted as the presence of molybdate in that position that would lock the catalytic site and allow the capture of transient complexes. However, for the sake of clarity and continuity, we treated all hydrolyzed nucleotides as ADP+Pi.

For dataset A, after successive rounds of 2D and 3D classifications, heterogeneous refinement, 3D variability analysis and final homogeneous and non-uniform refinements, we were able to isolate two distinctive classes: the RHC complex and a new RRHCC complex (Fig. 1B-C, Fig. S2E, processing branch I). The latter was further analyzed, and three subclasses were identified according to the visibility of CDC37.1-MD and the RAF1.2 C-lobe. Therefore, we characterized a class missing both as RHCC, which differed from the RHC particle in having a partially visible second molecule of CDC37 (Fig. S5). Another class was clearly a RRHCC particle, but CDC37.1-MD was not associated with HSP90 and consequently was not visible (RRHCC_c0). The remaining class, RRHCC with a visible CDC37.1-MD, was furthermore analyzed (3D variability analysis and 3D classifications) to resolve the heterogeneity of the mentioned domain (Fig. S2E branch II). All these classes displayed quite similar base structure with small differences, indicating that in their movement, the CDC37 MD acts as a rigid body (Fig. S8A). Despite the similarities, all these classes diverged from each other at the RAF1.2 αC helix extension and orientation (Fig. S7). When considering all variants, their respective movements can be described as rotations of the CDC37-MD over subunit A of the HSP90 dimer (Fig. S8B). Taking RRHCC_c4 as a reference, transformation matrices were calculated from its CDC37.1-MD to the positions of the equivalent domain in the rest of models, and the associated rotation axes were displayed for comparison (Fig. S8C). While a group of the axes concentrated, with similar orientations, near the HSP90-A portion close to the histidine pocket (RRHCC_c3, c6, a4, a6, Cloc), others (RRHCC_c1, c2 and c5) had quite different orientations. This could indicate that the different positions are more related with random jumps occurring while the interfaces adjust than to discrete gradual rotations around a common axis. We propose that this kind of vibration of CDC37.1-MD is used to constantly challenge the integrity of the αC helix of RAF1.2, preventing its extension into a fully folded helix and at the same time keeping a seed that eventually will trigger its complete folding, its dissociation from CDC37.1-MD and the reorganization of the N-lobe of RAF1.2.

The analysis of the interaction and dynamics of CDC37.1-MD with the partially folded N-lobe of RAF1.2 in the RRHCC complex revealed a structural specificity that checked for the presence of in the RHC particles (Fig. S2E branch IV). Class RHC_b was subjected to 3D classification and 3D variability analysis to provide four classes that described the heterogeneity in CDC37.1-MD. These classes, RHC_c1 to c4, represent different snapshots in the movement of the domain with respect of the rest of the complex and confirm the same type of interaction (Fig. S4).

For dataset B, following the same general procedure, we obtained three different classes. Two of them resembled previously discussed classes, RHC and RRHCC_c0-like particles, but we were able to isolate a distinctive class that, after structural analysis, was revealed to be an RHC particle containing p23 (RHCp23, Fig. 3, Fig. S2E branch V). Further variability analysis of the latter provided two final maps describing two states of the interaction of p23 with the RHC complex (Fig. 3). Both RHCp23 maps showed lower resolution values compared with all other RHC and RRHCC maps in this study at the HSP90 amino-terminal domains and specifically around the ATP binding sites (SI1-3). This could be interpreted as if the binding of p23 to the RHC complex injects stress into the structure and raises the conformational heterogeneity of the complex in contrast with the NTDs of HSP90. We hypothesize that this tension could be used to open the complex and release its cargo once both ATPs are hydrolyzed. We interpret the compositional difference between the two datasets as if they were sampled at different maturation states of the RAF1-CDC37-HSP90 complexes: dataset A represents the initial states containing RHC and RRHCC complexes, while dataset B represents a more aged population enriched in RRHCC_c0 and RHCp23.

Despite the preferential distribution of orientations showed by the RHC complexes^24^, the use of tilted images helped to prevent anisotropic maps (SI1-3 panels I-V). Some reconstructions presented relative low values for the validation metrics (cFAR and SCF^59^, mainly in RHCp23 classes), in contrast with the rest of the maps. Although there was some preference in the orientation of the complexes, the distribution was sufficient to prevent strong artifacts in the regions with highest resolution. We consider that the relatively low values of the sampling metrics in those specific cases are more related to the presence of low-resolution regions in quite confined regions due to extreme heterogeneity (i.e. p23, CDC37-MD), which differentially affects the directional signal evaluation. Global resolution of the non-uniform refined maps was estimated based on the gold standard FSC at 0.143 criterion between two half maps refined independently in CryoSPARC (SI1-3). Final maps were filtered according to the local resolution calculated in CryoSPARC with B-factor sharpening. In some figures, maps without sharpening were used for the sake of clarity (due to the large difference in detail between HSP90 and other more heterogeneous regions).

A comparison across all the structures showed that despite the different compositions of the complexes and the described movements of RAF1 and CDC37-MD, the overall scaffolding of HSP90 remained quite similar for all RHC/RRHCC/RHCp23 structures (Fig. S11). However, the alignment of the different structures in HSP90-A-NTD revealed small deviations that were amplified at the CTD of HSP90 (Fig, S11A). In this sense, RHC is the most different of the complexes, and the root of this difference seems to be the internal symmetry of the structures, with RRHCC being the most symmetric and RHC the least (Fig. S11B). This is not only reflected in the relative orientation of the HSP90-CTD but also in the NTDs of HSP90 (Fig. S11C), with differences that cannot be exclusively assigned to the state of the catalytic sites (compare RHC_b to RHCp23_c). The basis of the symmetry in the RRHCC complexes is evidently based in the binding of equivalent CDC37:RAF1 to both sides of HSP90 and can be extended to the RHCp23 complexes as p23 is occupying a similar position as the most N-terminal part of CDC37. All these diverse structural elements play simultaneously and in coordination to temporary sequester RAF1 and eventually produce a folded active kinase (SI4B).

### Model building and refinement

The previous model of the RHC particle^24^ was used as initial template for all the different complexes. AlphaFold models of CDC37 CTD and p23 were used for the RRHCC a4/a6 and RHCp23 complexes respectively ^60^. Rigid-body fitting, model modification, and energy minimization were performed using ChimeraX v1.10 ^61^ and Isolde ^62^, with additional editing in Coot v0.9 ^63^. Final refinement was performed using PHENIX v.2.0^64,65^. Statistics for the final maps and refinements are presented in Table S2. ChimeraX was used for analyzing maps and models and for generating figures.

### Phosphoproteomics

Digestion of purified samples of the isolated samples was performed using a protein aggregation capture (PAC)-based method with MagReSyn® Hydroxyl beads (ReSyn Biosciences). For each sample, 5eµg of purified protein was diluted in 50eµL of 70% acetonitrile (ACN). Hydroxyl beads were then added at a protein-to-bead ratio of 1:8. After a 10-minute incubation at room temperature (RT), the beads were washed twice with 100% ACN and once with 70% ethanol using a magnetic rack. Digestion was initiated by adding 50eµL of 50emM ammonium bicarbonate containing trypsin and Lys-C at protein-to-protease ratios of 1:100 and 1:250, respectively. Following overnight incubation at 37°C, the reaction was quenched by adding trifluoroacetic acid (TFA) to a final concentration of 1%. Evotip® Pure (Evosep) tips were washed with 20□µL of 100% ACN and centrifuged for 1 minute at 800 ×□g. Preconditioning was performed by adding 20□µL of 0.1% formic acid (FA) while soaking the tips in 100% isopropanol, followed by centrifugation for 1 minute at 800 ×□g. From each digested sample, 50eng of peptides was loaded onto the Evotip and centrifuged for 1 minute at 800 ×eg. Tip preparation was completed by sequentially adding 20eµL of 0.1% FA (1emin centrifugation), followed by 200eµL of 0.1% FA and a final centrifugation for 15 seconds. Samples were analyzed using an EV-1109 column (PepSep, 8ecm ×e150eµm, 1.5eµm beads) and an EV-1087 emitter (20□µm fused silica) on an Evosep One LC system. The column was coupled online via an EASY-Spray™ source to an Orbitrap Astral mass spectrometer (Thermo Scientific), operated using Xcalibur (tune version 4.0 or higher). The spray voltage was set to 1.8ekV, funnel RF level to 40, and capillary temperature to 275°C. Data-dependent acquisition (DDA) was performed with a 0.6-second cycle time, scanning from 250 to 1,560em/z, and a 5-second injection time. LC–MS/MS data were analyzed using SpectroMine (version 4.5.2), with searches performed against the Homo sapiens UniProt proteome (2023 release, 20,594 entries) supplemented with a database of common contaminants (381 entries). Phosphorylation of serine, threonine, and tyrosine (STY) was included as a variable modification, with post-translational modification (PTM) localization enabled. To compare phosphorylation sites in the RRHCC complex (this study) and the RHC complex^24^, mass spectrometry output files were analyzed in Rstudio using the following packages: dplyr (1.1.2)^66^, pheatmap (1.0.12)^67^, readxl (1.4.5)^68^, sva (3.48.0)^69^, tidyr (1.3.1)^70^, and ggplot2 (3.5.1)^71^. Output files from both datasets were imported using the readxl package. Data were then adjusted for batch effects using the ComBat function from the sva package. Raw intensities were subsequently log-transformed and visualized using the pheatmap package. The raw data of the phosphoproteomic analysis have been deposited into the PRIDE database with project accession code PXD065622.

### Mass spectrometry quantification

Proteins were solubilized in 2% SDS and 100 mM triethylammonium bicarbonate (TEAB, pH 7.55) and subsequently reduced and alkylated with 15 mM tris(2-carboxyethyl)phosphine (TCEP) and 25 mM chloroacetamide (CAA) for 1 h at 45°C in the dark. Samples were digested using the Protein Aggregation Capture (PAC) method^72^ with MagReSyn® Hydroxyl microparticles on a KingFisher instrument (Thermo Fisher Scientific). Digestion was carried out with a LysC/Trypsin mixture (Wako, Merck) at a 1:50 protein-to-enzyme ratio for 16 h at 37°C in 50 mM TEAB (pH 8.0). Resulting digests were acidified with formic acid (FA) to a final concentration of 0.5% (v/v).

LC-MS/MS analyses were performed on a Vanquish™ Neo HPLC system coupled either to an Orbitrap Astral mass spectrometer (heterodimer/homodimer samples) or to an Orbitrap Exploris 480 (acidic patch and RAF1 loop mutants) (Thermo Fisher Scientific). Peptides were loaded onto a trap column (PepMap™ Neo, 5 µm C18, 300 µm × 5 mm) and separated on either an Aurora Elite TS analytical column (C18, 1.7 µm, 75 µm × 25 mm) or an EASY-Spray™ PepMap™ Neo column (C18, 2 µm, 75 µm × 500 mm), both operated at 50 °C. Buffer A consisted of 0.1% FA and Buffer B of 100% acetonitrile (ACN). Peptides were eluted using a 2–45% Buffer B gradient over 50–60 min at a flow rate of 400 nL/min, followed by a 7-min wash at 98% Buffer B. Ionization was performed using an EASY-Spray source at 1.8 kV (heterodimers/homodimers) or 1.5 kV (acidic patch and RAF1 loop mutants). The ion transfer tube temperature was set to 300°C.

MS acquisition was performed in data-independent acquisition (DIA) mode every 0.6 s for heterodimer/homodimer samples or every 75 scans for mutant samples. Precursor scan ranges were 390–380 m/z (heterodimers/homodimers) and 390–1000 m/z (mutants). Injection times ranged from 3 to 22 ms. Orbitrap MS1 resolution was set to 240,000 for heterodimers/homodimers and 60,000 for acidic patch/RAF1 loop mutants. MS/MS scans were acquired at 80,000 or 15,000 resolution, respectively, depending on the instrument.

The raw data for RAF1 and acidic patch mutants were analyzed using DIA-NN 1.9.2 in library-free mode against a custom database containing the overexpressed constructs and the human UniProtKB/Swiss-Prot proteome (one sequence per gene, 20,614 entries). The precursor m/z range was set to 390–1010. Protein abundances were calculated by summing precursor intensities reported in report.tsv, filtered for Lib.Q.Value and Lib.PG.Q.Value < 0.01. Only unique peptides were used for quantification. Intensities were normalized to the total protein intensity per sample. Isoform-specific CDC37 peptides were excluded to ensure consistent comparisons across conditions. For RAF1 mutants, only peptides shared by all mutated forms were used for quantification.

The raw DIA files from heterodimers/homodimers were processed using DIA-NN 1.9.2 in library-free mode against the same human database described above. The precursor m/z range was set to 379–981. Quantification was performed by summing unique precursor intensities after applying quality filters (Lib.Q.Value and Lib.PG.Q.Value < 0.01). Isoform-specific B-RAF peptides were excluded. Intensities from purified samples were normalized to the corresponding whole-cell lysate to account for input variability.

### Site-directed mutagenesis and pulldowns from RAF1 and CDC37 mutants

All plasmids containing RAF1 mutants (H402R, H402D, H402A, N404A, F408R, F408E, R401A) were generated from pcDNA3-RAF1-Strep and pcDNA3-CDC37-Strep, respectively, using the QuikChange™ Lightning Site-Directed Mutagenesis Kit (Agilent Technologies) following the manufacturer’s instructions and using the primers listed in Supp. Table 3. To study the effect of RAF1 mutants on RAF1 stability, we followed a slightly modified version of the protocol described previously^24^. Briefly, 20 mL of Expi293F cells was transiently transfected with: A.-For RAF1 mutant analysis: 5 μg of pcDNA3-HSP90-HA, 5 μg of pCMV-CDC37-myc-DDK, and 10 μg of pcDNA3-RAF1-Strep (wild-type or mutant). B.-For CDC37 mutant analysis: 5 μg of pcDNA3-HSP90-HA, 10 μg of pcDNA3-RAF1-V5, and 5 μg of pcDNA3-CDC37-Strep (wild-type or mutant).

Forty-eight hours post-transfection, cells were harvested and lysed in buffer (20 mM Tris-HCl pH 7.5, 150 mM NaCl, 10 mM MgCle, 10 mM KCl, 20 mM NaeMoOe, and 0.1% Triton X-100) supplemented with cOmplete™ Mini Protease Inhibitor Cocktail, Phosphatase Inhibitor Cocktails 2 and 3, and benzonase nuclease. Thirty milligrams of whole-cell lysate was incubated with 200 μL of Strep-Tactin® XT 4Flow High-Capacity resin for 10 minutes at 4°C. After washing with 12 CV of the same buffer (without supplements), proteins were eluted with biotin. One-tenth of each eluate was analyzed by western blotting, and RAF1 levels were quantified by mass spectrometry analysis.

### Pulldowns from CDC37-Strep and his-tagged kinases

To assess CDK4 dimer formation in chaperone complex, Expi293F cells were transfected with CDC37-StrepTag, Halo-KRAS^G12V^, V5-and HisTagged RAF1 or CDK4. After Strep purification as previously described, eluates were purified using 40 µL HisPur™ Ni-NTA magnetic beads (Thermo Fisher Scientific). Protein elution was achieved by high-concentrated imidazole buffer and V5-proteins eluates were analyzed by immunoblotting.

### Western blot analysis and antibodies

Twenty micrograms of whole-cell lysate or one-tenth of the eluted samples after the double affinity purifications was separated by SDS-PAGE using NuPAGE™ 4–12% Bis-Tris Midi Gels (ThermoFisher Scientific) and transferred to nitrocellulose blotting membranes (Cytiva Life Sciences). Following blocking with 5% BSA in TBS-T for 1 hour at RT, the membranes were incubated with the following primary antibodies: anti-HSP90 (Santa Cruz, sc-13119; 1:1,000), anti-RAF1 (BD, 610151; 1:1,000), anti-CDC37 (Santa Cruz, sc-13129; 1:1,000), anti-pan-Ras (Calbiochem, OP40; 1:1,000), anti-V5-tag (ThermoFisher Scientific, R960-25; 1:1,000), anti-StrepTagII (ThermoFisher Scientific, MAS-37747; 1:500), anti-Vinculin (Sigma, V9131; 1:5,000), anti-BRAF (Santa Cruz, sc-5284; 1:1,000), anti-ARAF (Cell Signaling, 4432; 1:500), anti-phospho-MEK1/2 (Ser217/221) (Cell Signaling, 9121; 1:250), anti-MEK1 (Santa Cruz, sc-6250; 1:1,000), anti-ERK1/2 (BD, 554100/610103; 1:1,000), anti-phospho-ERK1/2 (Cell Signaling, 9101; 1:250) and anti-CDKN2A/p16-INK4A (Proteintech, 10883-1-AP; 1:500). Detection was performed using either HRP-conjugated secondary antibodies and Amersham™ ECL Western Blotting Detection Reagent (Cytiva Life Sciences), or fluorescent dye-conjugated secondary antibodies and visualized using the Odyssey CLx imaging system (LI-COR Biosciences).

### Heterodimer and homodimer formation assay

For pull-down experiments, full-length human ARAF, BRAF, CDK4, and a fusion protein comprising RBD-CRD(RAF1)-CDK4 fused to either a StrepTagII or a V5-tag were cloned into the pcDNA3.1 vector using the In-Fusion HD EcoDry Cloning Kit (Takara Bio). The BRAF^D594A^ and BRAF^V600E^ mutants were generated using the QuickChange Lightning Site-Directed Mutagenesis Kit. To assess heterodimer formation, 20 mL of Expi293F cells was transfected with 4 μg of pFN21a-Halo-KRAS^G12V^, 8 μg of pCMV-CDC37-myc-DDK, 4 μg of pcDNA3-RAF1-Strep, and 4 μg of the desired RAF family member or CDK4 fused to a V5-tag. As a control, 8 μg of pcDNA3-RAF1-V5 was transfected under the same conditions. For homodimer detection, the same strategy was used, co-transfecting 4 μg of the Strep-tagged construct and 4 μg of the corresponding V5-tagged version of the same protein. The expression and pull-down protocol were the same as that used for the RRHCC complex, scaled accordingly. Briefly, 48 hours post-transfection, cells were harvested and lysed in KRC buffer as previously described, and 40 mg of lysate was incubated with 200 μL of Strep-Tactin® XT 4Flow High Capacity resin, pre-equilibrated in KRC buffer, for 10 minutes at 4°C. After washing with 12 CV of KRC buffer, bound proteins were eluted with 5 mM biotin. The eluate was incubated with 200 μL of HaloLink™ resin for 1 hour at 4°C and washed with 12 CV of KRC buffer. Proteins were eluted with 7.5 U of Halo-TEV protease. One-tenth of each eluate was analyzed by SDS-PAGE followed by western blotting using an anti-V5 antibody to detect heterodimers or homodimers. Protein levels were further quantified by mass spectrometry. All experiments were performed in duplicate.

### Phosphorylation in RAFless MEFs

To generate viral particles encoding the different RAF isoforms, we constructed several expression plasmids. The retroviral vector pBABE-hygro-ARAF was generated using the In-Fusion HD EcoDry Cloning Kit. Retroviral vectors pBABE-hygro-BRAF WT, BRAF^D594A^, and BRAF^V600E^ were already available in our lab. For heterodimer studies involving RAF1, the lentiviral vector pLVX-puro-RAF1 described previously^24^ was used. Retroviral and lentiviral particles were produced by transient transfection of HEK293T cells using standard packaging plasmids. Resulting virus-containing supernatants were used to infect immortalized *ARaf^lox/lox^*; *BRaf^lox/lox^*; *Raf1^lox/lox^* (RAFlox) mouse embryonic fibroblasts (MEFs). Following infection and antibiotic selection with the appropriate resistance markers, cells were transduced with adenoviral particles encoding Cre recombinase to induce genetic ablation of endogenous *RAF* (RAFless). Seventy-two hours post-infection, cells were harvested, and MEK1 phosphorylation was analyzed by Western blotting.

### RAFless proliferation assay

MEFs were generated as described above. To evaluate their proliferative capacity, cells were infected with retroviral or lentiviral particles encoding the indicated RAF isoforms, with or without a short hairpin RNA (shRNA) targeting *p16^INK4a^* to prevent the onset of cellular senescence, as previously reported ^54^. After antibiotic selection, 10,000 infected cells were seeded per p100 plate in DMEM supplemented with 10% (v/v) FBS and maintained for 10–14 days at 37°C in a humidified atmosphere containing 5% COe. The culture medium was replaced every 3–4 days. At the endpoint, colonies were fixed with 4% paraformaldehyde for 15 min at RT, washed twice with PBS, and stained with 0.5% (w/v) crystal violet (Sigma). Excess dye was removed by rinsing with distilled water, and plates were air-dried. For quantification, bound crystal violet was solubilized, and absorbance was measured spectrophotometrically. Data were normalized to RAFless control cells.

## SUPPLEMENTARY MATERIALS INDEX

**Figs. S1 to S11**

**Supporting Information Legends 1-4**

**Table S1. Phosphorylation of RAF1-chaperone complexes Table S2. CryoEM data collection and refinement**

**Table S3. Oligonucleotides, cell lines, etc.**

**Figure S1. Characterization of the RAF1-chaperone preparation used for structure determination and phosphoproteomics analysis of the purified RRHCC sample and comparison with the RHC sample, related to STAR Methods and Table S1. (A)** Coomassie-stained SDS-PAGE analysis of the RRHCC sample following two-steps affinity chromatography (StrepTag-RAF1 and HaloTag-KRAS purification). **(B)** Western blot confirming the identity of selected protein bands observed in the Coomassie-stained gel shown in (A). **(C)** Elution profile of the RRHCC sample obtained by size exclusion chromatography (SEC) using a Superdex 200 column. **(D)** Coomassie-stained SDS-PAGE analysis of SEC-eluted fractions. **(E)** Mass spectrometry analysis of the affinity-purified extract containing the RAF1-chaperone complexes, along with sub-stoichiometric amounts of co-purifying 14-3-3 proteins. The plot displays mean iBAQ protein levels from a representative experiment. **(F)** Domain architecture of RAF1, including the phosphosites found in our analysis of the RRHCC sample compared with the RHC sample ^24^. **(G)** Table including the phosphosites found in the RHC and RRHCC samples. **(H)** Heatmap depicting the differential phosphorylation intensity of 16 key residues in RAF1 (Table S1).

**Figure S2. Cryo-EM data and processing workflow, related to** Figure 1, 2**, 3, 4, Table S2. (A)** Representative motion-corrected cryo-EM micrograph of the RHC particles in vitreous ice imaged with a Titan Krios Falcon 4i direct detector camera (-2.3 μm defocus). **(B)** Characteristic reference-free 2D averages of the RHC particles. The scale bar represents a length of 100 Å. **(C)** Similar 2D averages of the RRHCC particles, with the characteristic extra CDC37 and RAF1 subunits indicated (white arrows and triangle respectively). **(D)** Similar 2D classes of the RHCp23 particle. The asterisk indicates the additional density corresponding with the p23 subunit. **(E)** Workflow diagram of the different strategies followed in the cryo-EM data processing and map refinement of the RHC/RRHCC/RHCp23 complexes.

**Figure S3. Structural details of the RRHCC complex, related to 1, 2, 4. (A)** Overall view of the RRHCC assembly. The HSP90 monomers are colored in transparent grey to show the entangled assembly between the RAF1 and CDC37 molecules around the HSP90 closed conformation. **(B)** Zoomed view of the ATP sites of the RRHCC_a6 complex. **(C)** Detailed view of pS13 in the two CDC37 molecules and its interaction in each side of the clamp with the corresponding HSP90 monomers. **(D)** View of the segment of the RAF1.1 protomer threading through the HSP90 lumen. **(E)** Local map showing the interaction of the section of the RAF1.2 histidine anchor with HSP90 and CDC37. The zoom depicts the H402 inside the cavity formed in the MD-CTD of one of the HSP90 monomers within the coulombic density of the region.

**Figure S4. Structures of the RHC and conformational landscape, related to** Figure 1, 2**, 4. (A)** Cryo-EM maps of the four RHC classes (c1-c4) resolved by variability analysis of CDC37.1-MD, colored according to their protein composition. **(B)** Ribbon representation of the same classes. Arrows indicate the relative movement of the CDC37.1-MD of classes c2-c4 in comparison to class c1. **(C)** Detail view of CDC37.1-MD interacting with the αC helix of RAF1.1 (local resolution filtered map depicted as a transparent grey surface). **(D)** Movement of CDC37.1-MD of the RHC classes shown in three perpendicular views. The direction of the movement (black arrow) is derived from the composition of the nucleotide binding site of HSP90-A. **(E)** Zoomed view of the ATP sites of the four RHC classes.

**Figure S5. Structural details of the RHCC complex, related to** Figure 1, 2**, 4. (A-B)** Ribbon diagram of the RHCC complex including the additional molecule of CDC37 in the coulombic density (pink). The zoom shows a detailed view of the interaction region allowing the modeling of the β-strand of the second molecule of CDC37 building the antiparallel β-sheet with HSP90. **(C)** Zoomed view of the ATP sites of the RHCC classes. **(D-E)** Zoom of the pS13 in both CDC37 protomers of the RHCC complex, facilitating the assembly of the co-chaperone with HSP90.

**Figure S6. Comparison and structural alignment of the RAF1 kinase molecules visualized in the RRHCC complex, related to** Figure 1, 2**. (A-B)** Structural comparison of the two kinase domains of RAF1 found in the RRHCC complex (dark and light green) depicting the different folding stages and the RAF1 folded kinase domain (PDB:8CPD) (white). The orange-red segment corresponds to the non-visualized section of RAF1 that folds into β1 to β3. The connecting loops and the first turn of the αC helix in the kinase domain structure are unfolded in the RHC complex and in RAF1.1 in the RRHCC complex (light green) but are folded in RAF1.2 in the RRHCC complex (dark green). **(C)** Amino acid sequences of human RAF1, ARAF, BRAF and CDK4 were aligned using Clustal Omega (http://www.ebi.ac.uk/Tools/msa/clustalo). The figure was prepared with ESPript (http://espript.ibcp.fr). Residue numbers are labelled according to the RAF1 sequence. Similar residues are shown in red and identical residues in white over red background. Secondary structures including α helix and β sheets for RAF1 are indicated. The upper bar and lower bar in the alignment are colored according to each RAF1 molecule. The orange-red sector corresponds to the regions not visualized in the kinase domain.

**Figure S7. Conformational heterogeneity of the** α**C helix of RAF1.2 in the CDC37.1-MD/HSP90-A mold, related to** Figure 1, 2**. (A)** The analysis of the αC helix of RAF1 in several RRHCC classes shows heterogeneity in what can be identified as different states of folding, with a variable span of the alpha helix. Upper panels show the interaction of the trapped part of the N-lobe of RAF1 (green, with its αC helix in light sea green and H402 in lime) between CDC37.1-MD (orange) and HSP90-A (blue). The middle panels depict the corresponding maps. The lower panels show exclusively the densities and models of the trapped N-lobe. **(B)** Different view after 60° rotation.

**Figure S8. Detailed conformational landscape in the mold in the RRHCC complex and interaction of CDC37.1-CTD with RAF1.1 in the region where the RAF1-14-3-3 complex associates with MEK, related to** Figure 1, 2**. (A)** Three perpendicular views of the RRHCC classes indicated in the inset, where the different structures are aligned according to their CDC37-MD. **(B)** Same as (A) but with all the structures aligned according to HSP90-A and hence showing the relative movement of CDC37.1-MD. **(C)** Rotation axes summarizing the movement from the initial position of RRHCC c4, taken as reference, to all other structures. The values of the rotation associated with each axis and the translation along them are indicated in the inset. **(D)** Comparison of two RRHCC classes with the CDC37.1 carboxy-terminal domains oriented towards the RAF1.1 C-lobe, in two views. **(E)** Superposition of RRHCC_a6 and a RAF1:MEK1 complex (PDB:9MMP, CRAF/MEK1/14-3-3 complex), fitted in RAF1, depicting how this orientation of the CDC37.1-CTD can block the premature association of RAF1 with its substrate. **(F)** Detailed views showing the associated Cryo-EM densities of both RRHCC_a4 and RRHCC_a6 classes. **(G)** Tentative ribbon model of the possible organization of the RAF1:CDC37 complex (RC), based in the different contacts between these two components identified in the RRHCC complex (Fig. 1, 2, S8D-F). **(H)** Depiction of how the previously described RC complex could be loaded into HSP90 to produce the RHC and RRHCC complexes. The latter could be achieved by the successive addition of two RC or by the incorporation of a 2xRC complex (in which there is a domain swapping of the αC helix of RAF1 of two RC; and its loading requires breaking the interaction of the αC helix of RAF1.1 with CDC37.2-MD and capture of the RAF1.1 N-lobe by HSP90, while maintaining all the rest of interactions).

**Figure S9. Controls for RAF1, CDK4 and RAF family homo-and heterodimerization, related to** Figure 5**. (A, D)** Western blot analysis of input lysates and pull-down samples from the experiments described in Figure 5A and Figure 5E, respectively. **(B)** Schematic representation of the experimental workflow used to detect CDK4 homodimers by sequential Strep-Tag and HisTag purifications from clarified cell lysates. **(C)** Western blot analysis of input lysates and pull-down samples from the experiment described in (B). Solid arrowheads indicate the migration positions of the relevant proteins. Tagged proteins were detected using tag-specific antibodies. Abbreviations: St, StrepTagII; HT, HaloTag; His, HisTag. Complex notations: KXStXV5C = HaloTag-KRASG12V + X–StrepTagII + X–V5 + CDC37–Flag, where X represents: R= RAF1, A = ARAF, B = BRAF, B* = AFD594A, B** = BRAFV600E, C = CDK4. KXHisXV5C = HaloTag-KRASG12V + X–HisTag + X–V5 + CDC37–StrepTagII, where X represents: R = RAF1; C = CDK4. RStXV5C=HaloTag-KRASG12V + RAF1–StrepTagII + X–V5 + CDC37–Flag, where X represents: A =ARAF, B = BRAF, B* = BRAFD594A, B** = BRAFV600E.

**Figure S10. Ectopic expression of RAF dimers and proliferation levels in RAFless MEFs, related to** Figure 6**. (A)** Western blot analysis of lysates from the experiment described in Figure 6B without or **(B)** after co-infection with shRNA targeting p16INK4a. **(C)** Quantification of two independent experiments after dissolving the crystal violet and measuring absorbance from the experiment shown in Figure 6B. Light purple bars correspond to samples without p16 knockdown, and dark purple bars correspond to samples expressing the p16-shRNA. Absorbance values are shown relative to RAFless cells.

**Figure S11. Overall structural analysis of the RHC/RRHCC/RHCp23 complexes, related to 1, 2, 3, 4 and STAR Methods. (A)** Comparison of pairs of structures fitted by the NTD of HSP90-A. The first member of each couple is colored in gray, the second following the pattern in Fig. 1. Black arrows depict differences in the positioning of the HSP90-CTD. **(B)** The fitting of the HSP90-NTD of subunits B into A displays the internal symmetry of the different complexes. Small black arrows at the HSP90-CTD illustrate the divergences from perfect symmetry, as they amplify farther away. **(C)** Detail views of the HSP90-NTD not being fitted, with only RHC revealing major differences.

**Supporting Information 1. Experimental map analysis and validation, related to** Figure 1, 2**, 3, 4. (A-F)** Local resolution and validation of the 3D reconstruction of RHC_b and RRHCC classes. The cryo-EM density map in each case is shown in two opposite views and colored based on its local-resolution estimation (see common color key in a). A third panel is included, showing a cut-away view of the map of each complex. Validation analysis of the EM maps: (I) Gold-standard FSC curves with the estimated global resolution; (II) Conical FSC summary plot and cFAR value; (III) Viewing direction distribution plot; (IV) Relative signal amount vs. viewing direction plot and its representation in 3D; (V) Fourier sampling plots and SCF value.

**Supporting Information 2. Experimental map analysis and validation (continuation), related to** Figure 1, 2**, 3, 4. (A-F)** Local resolution and validation of the 3D maps of the rest of the RRHCC classes and the local map RRHCC_Cloc. Same panels as in SI1. Local-resolution color key is shared by A-E (F has its own key).

**Supporting Information 3. Experimental map analysis and validation (continuation), related to** Figure 1, 2**, 3, 4. (A-F).** Cryo-EM map analysis and validation (continuation). (A-F) Local resolution and validation of the 3D maps of the four RHC classes and the two RHCp23 classes. Same panels as in SI1.

**Supporting Information 4. Nucleotide occupancies in the HSP90 ATP binding pockets in the RAF1-chaperone complexes, related to** Figure 1, 2**, 3, 4, 7, SD4 (A)** The figure compiles together with Fig. 4 all the nucleotide pockets for the rest of structures described in this manuscript. See also Fig. 4, S3B, S4E and S5C. **(B)** Schematic illustrating the events in RAF1 maturation. For RAF1 to achieve its native folded state, the assembled αC helix must be released, and the HSP90 clamp must transition to an open conformation. These processes are interdependent and governed by the stability and extension of the αC helix bound to the CDC37-MD and ATP hydrolysis by HSP90. Failure in either step leads to incomplete RAF1 release or an unfolded kinase. The composition and conformational landscape of the RAF1–chaperone complexes collectively modulate these events.

