## Supplementary material for "The HSP90–CDC37 Chaperone System Orchestrates RAF1 Kinase Activation Through a Pre-Dimerization Mechanism": Fig. S1

### Figure S1

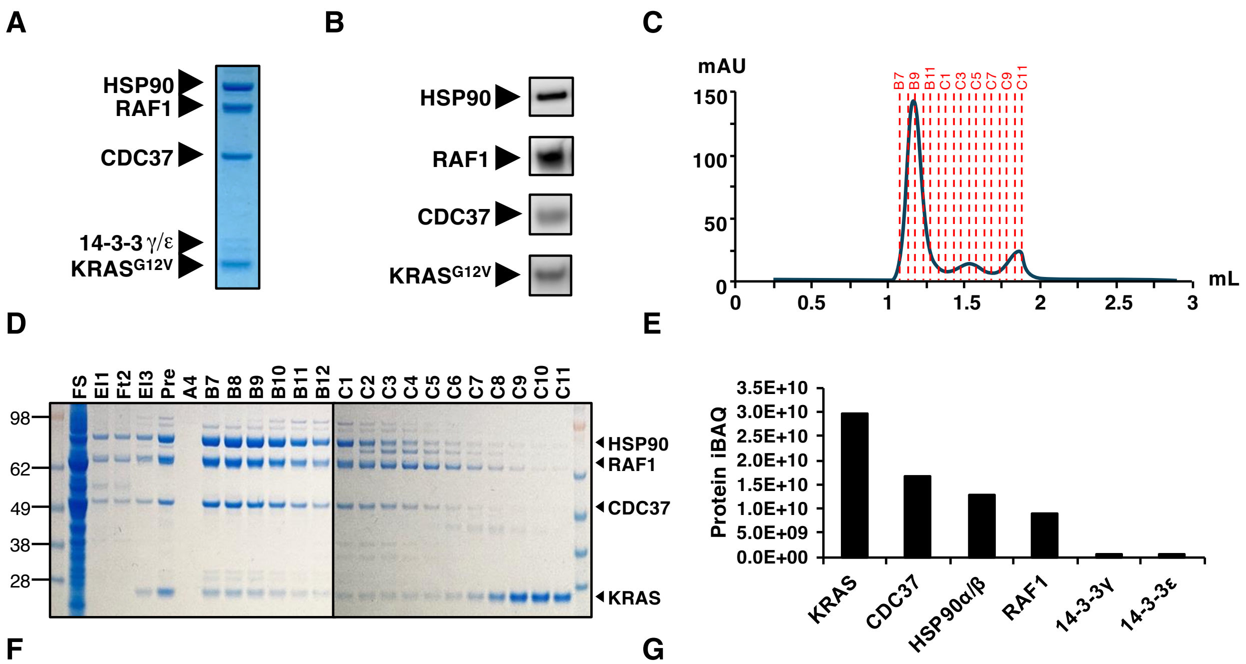


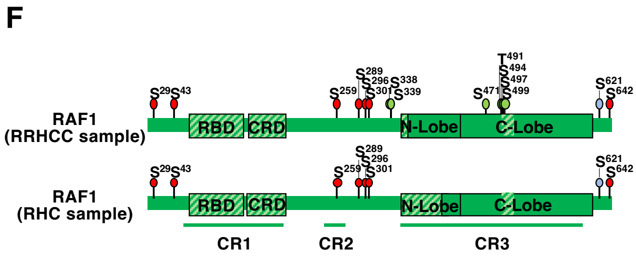


G

| | RHC sample[1] | RRHCC sample | | |
|---|---|---|---|---|
| | Phosphosite | Phosphosite | Site Probability | Flanking Region |
| CDC37 | Y4 | Y4 | 1 | MVDYSVWDHIE |
| | - | S5 | 1 | MVDYSVWDHIEV |
| | S13 | S13 | 1 | VWDHIEVSDDEDETH |
| | - | T19 | 1 | VSDDEDETHPNIDTA |
| | S27 | - | - | - |
| | - | Y298 | 1 | GLDPVEVYESLPEEL |
| | - | S300 | 1 | DPVEVYESLPEELQK |
| | - | T370 | 1 | LLEAVPKTGDEKDVS |
| HSP90AB1 | S226 | S226 | 1 | KEREKEISDDEAEEE |
| | S255 | S255 | 1 | PKIEDVGSDEEDDSG |
| | - | S261 | 1 | GSDEEDDSGKDKKKK |
| HSP90AA1 | - | T7 | 1 | MPEETQTQDQPMEE |
| | - | S231 | 1 | KERDKEVSDDEAEEK |
| | - | S263 | 1 | PEIEDVGSDEEEEKK |
| | - | S476 | 1 | ASGDEMVSLKDYCTR |
| HSP90AA2P | - | Y216 | 1 | KHSQLIGYPITLFVE |
| HSP90AA4P | - | S168 | 1 | KKNKIKLSARRDLIM |
| HSP90AA5P | - | S89 | 1 | PEIEDVGSDEEEEKK |
| RAF 1 | S29 | - | - | - |
| | S43 | S43 | 1 | FGYQRRASDDGKLTD |
| | - | S227* | 0.999 | SVSRMPVSSQHRYST |
| | S228 | S228 | 0.999 | VSRMPVSSQHRYSTP |
| | S233 | S233 | 1 | VSSQHRYSTPHAFTF |
| | - | Y232* | 1 | PVSSQHRYSTPHAFT |
| | T234 | T234 | 0.999 | SSQHRYSTPHAFTFN |
| | T239 | T239 | 1 | YSTPHAFTFNTSSPS |
| | - | T242* | 0.999 | PHAFTFNTSSPSSEG |
| | S244 | S244 | 1 | AFTFNTSSPSSEGSL |
| | - | S246* | 0.999 | TFNTSSPSSEGSLSQ |
| | - | S250* | 0.999 | SSPSSEGSLSQRQRS |
| | - | S252* | 1 | PSSEGSLSQRQRSTS |
| | S257 | S257 | 0.999 | SLSQRQRSTSTPNVH |
| | - | T258* | 0.999 | LSQRQRSTSTPNVHM |
| | S259 | S259 | 1 | SQRQRSTSTPNVHMV |
| | - | T260* | 0.999 | QRQRSTSTPNVHMVS |
| | - | S283* | 1 | MIEDAIRSHSESASP |
| | - | S285* | 1 | EDAIRSHSESASPSA |
| | - | S287* | 1 | AIRSHSESASPSALS |
| | S289 | S289 | 1 | RSHSESASPSALSSS |
| | S291 | S291 | 1 | HSESASPSALSSSPN |
| | - | S294* | 0.999 | SASPSALSSSPNNLS |
| | S295 | S295 | 0.999 | ASPSALSSSPNNLSP |
| | S296 | S296 | 1 | SPSALSSSPNNLSPT |
| | S301 | S301 | 1 | SSSPNNLSPTGWSQP |
| | - | T303* | 1 | SPNNLSPTGWSQPKT |
| | - | S306* | 1 | NLSPTGWSQPKTPVP |
| | - | T310* | 1 | TGWSQPKTPVPAQRE |
| | - | S338 | 1 | RPRGQRDSSYYWEIE |
| | - | S339 | 0.999 | PRGQRDSSYYWEIEA |
| | - | S357* | 1 | MLSTRIGSGSFGTVY |
| | - | S359* | 1 | STRIGSGSFGTVYKG |
| | - | T362* | 1 | IGSGSFGTVYKGKWH |
| | - | S471 | 1 | IIHRDMKSNNIFLHE |
| | - | T491 | 1 | IGDFGLATVKSRWSG |
| | - | S494 | 0.805 | FGLATVKSRWSGSQQ |
| | - | S497 | 1 | ATVKSRWSGSQQVEQ |
| | - | S499 | 0.999 | VKSRWSGSQQVEQPT |
| | - | T506* | 0.999 | SQQVEQPTGSVLWMA |
| | - | S508* | 0.999 | QVEQPTGSVLWMAPE |
| | - | S567* | 1 | MVGRGYASPDLSKLY |
| | S604 | S604 | 1 | PLFPQILSSIELLQH |
| | S605 | - | - | - |
| | - | S612* | 1 | SIELLQHSLPKINRS |
| | - | S619* | 0.756 | SLPKINRSASEPSLH |
| | S621 | S621 | 1 | PKINRSASEPSLHRA |
| | - | S624* | 0.956 | NRSASEPSLHRAAHT |
| | S642 | - | - | - |

[1] García-Alonso et al., Mol Cell. 2022

* New phosphosite identified in this work

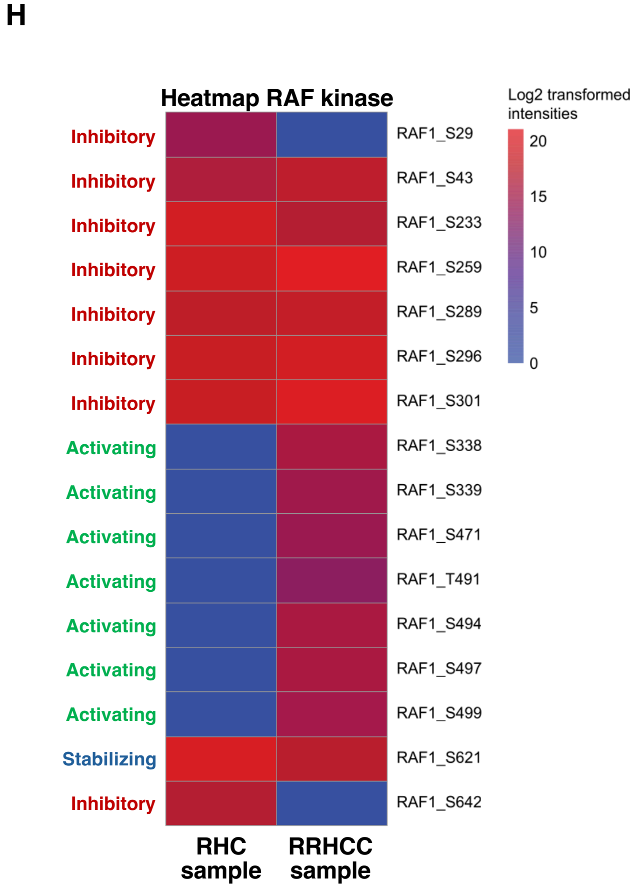
