## Supplementary figures and images for "The HSP90–CDC37 Chaperone System Orchestrates RAF1 Kinase Activation Through a Pre-Dimerization Mechanism"

### Fig. S2

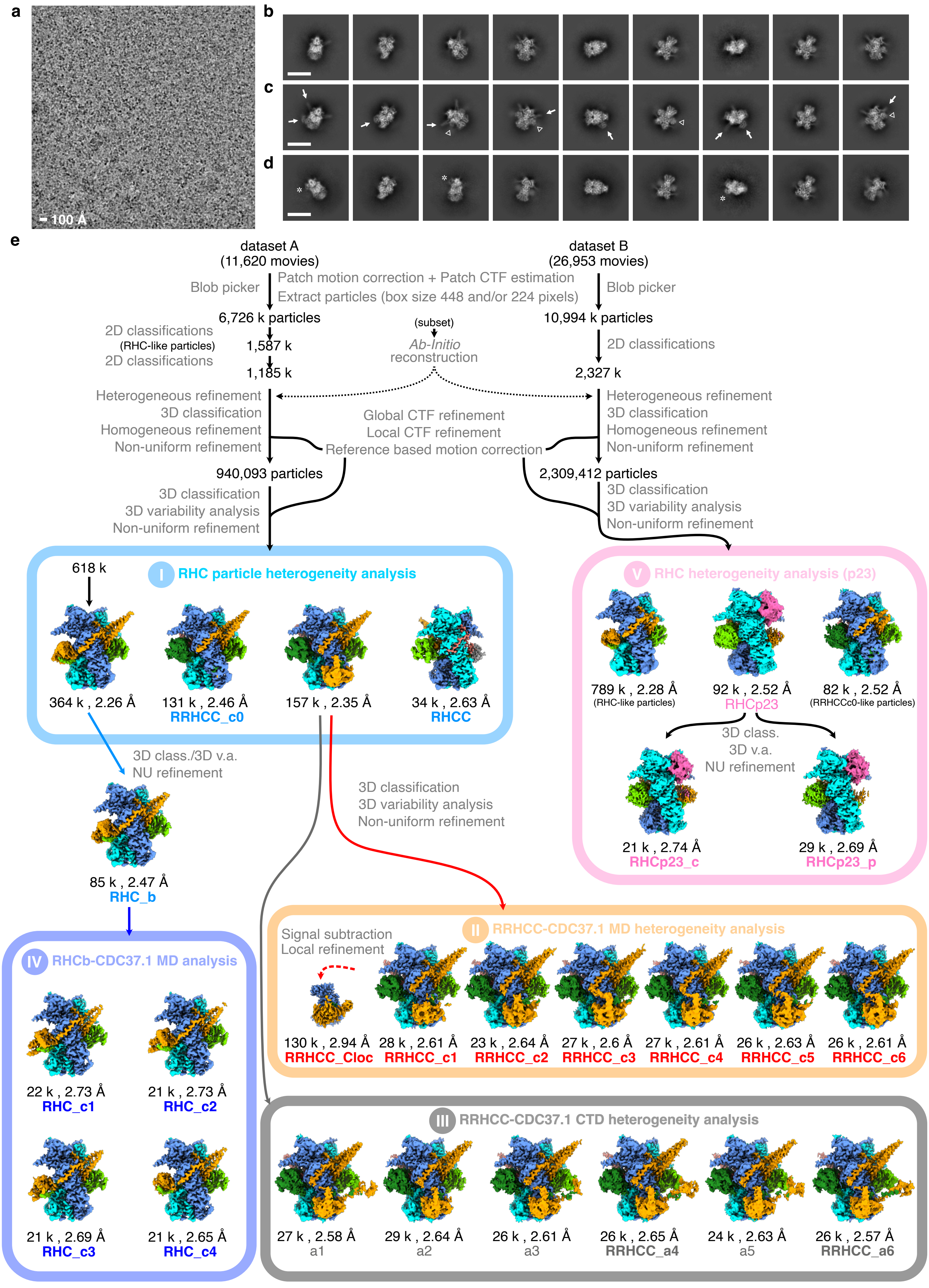

### Fig. S3

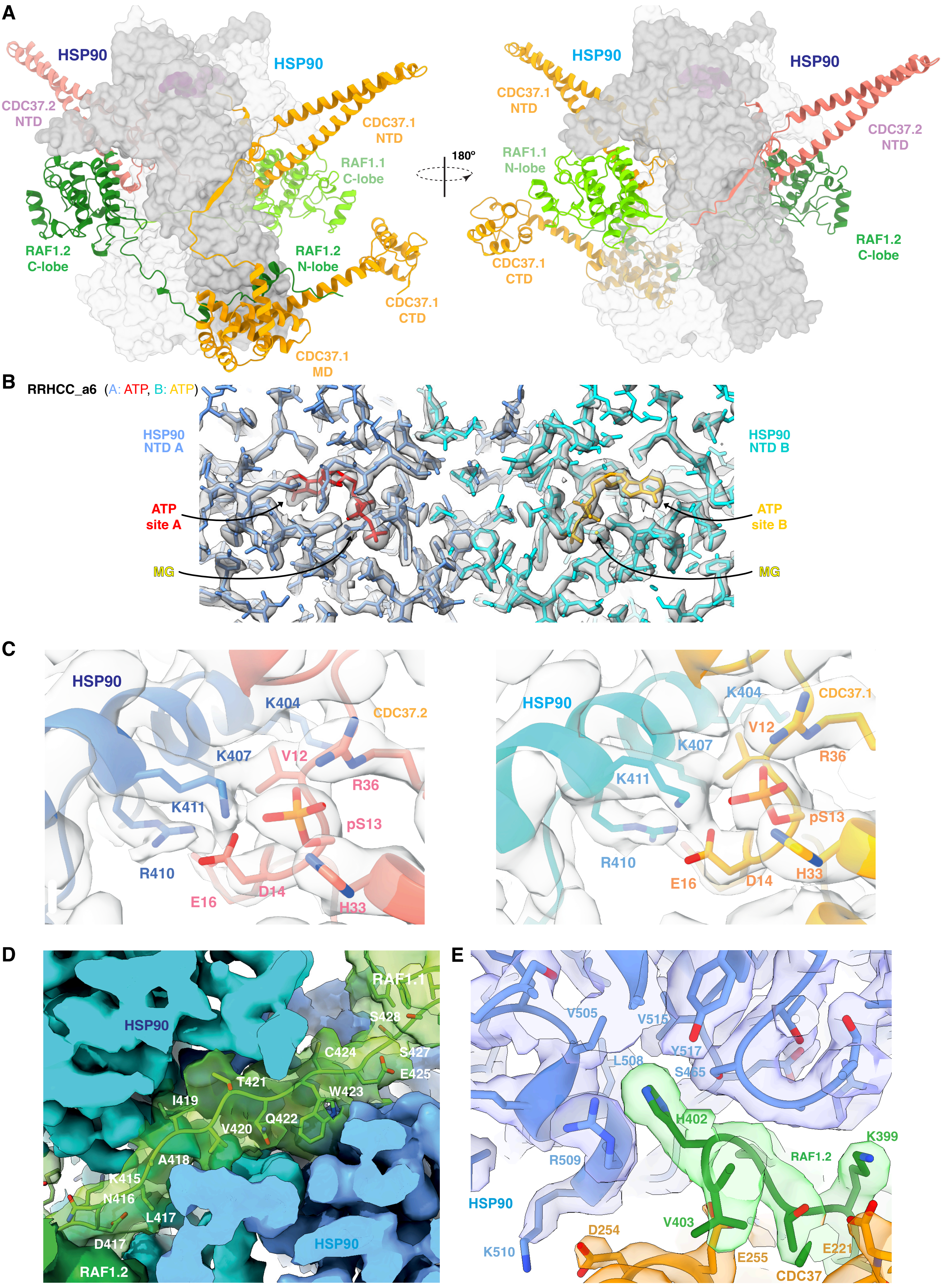

### Fig. S4

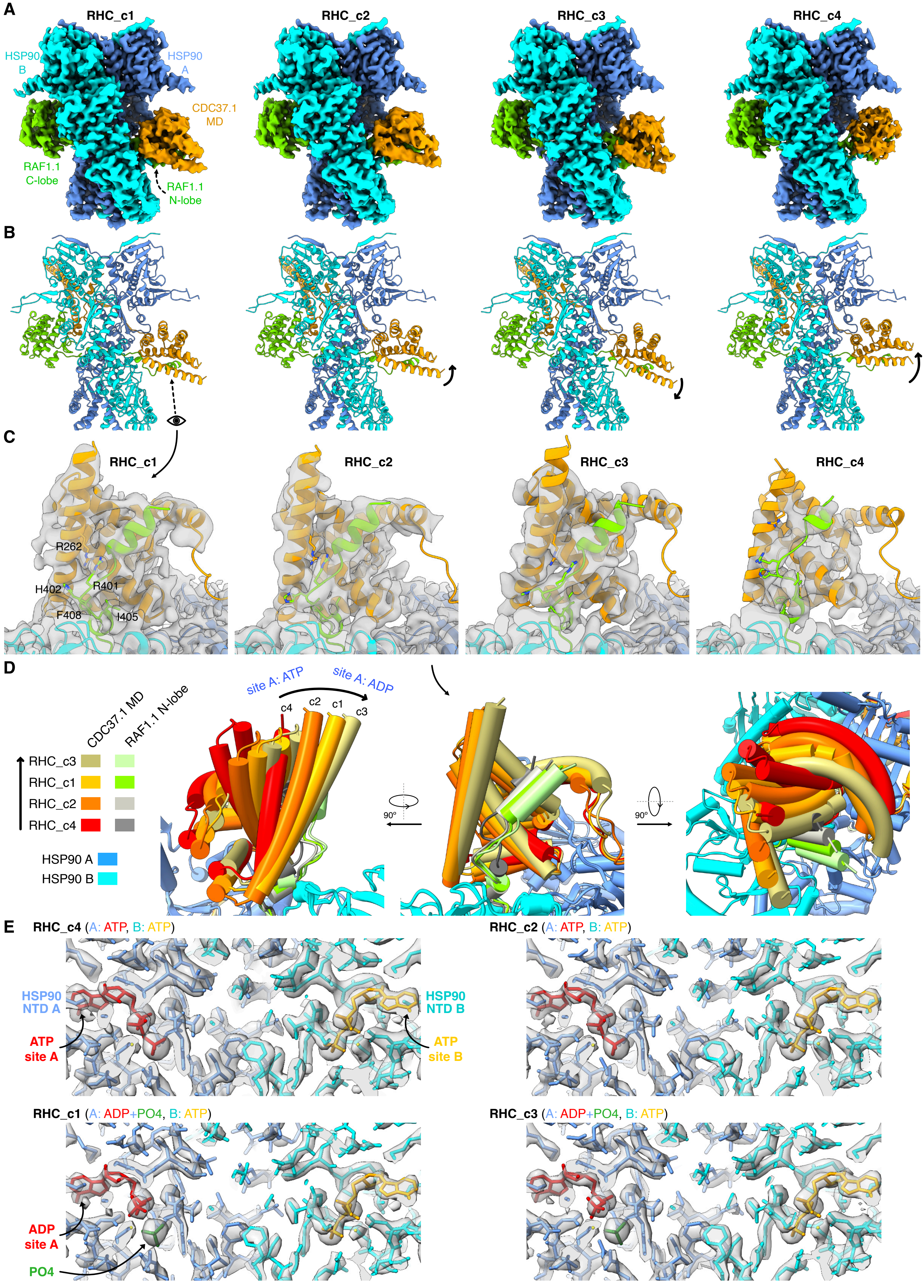

### Fig. S5

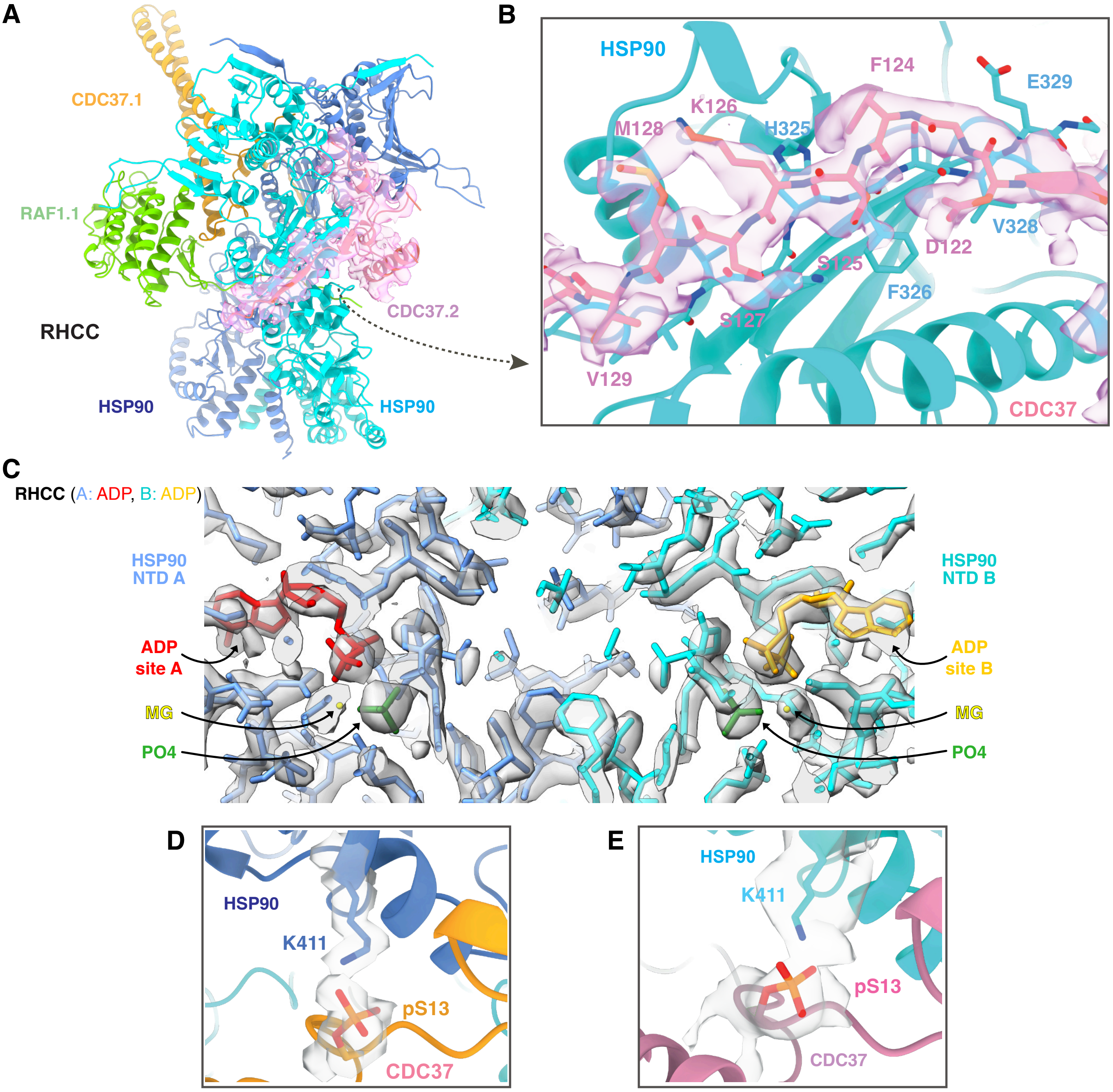

### Fig. S6

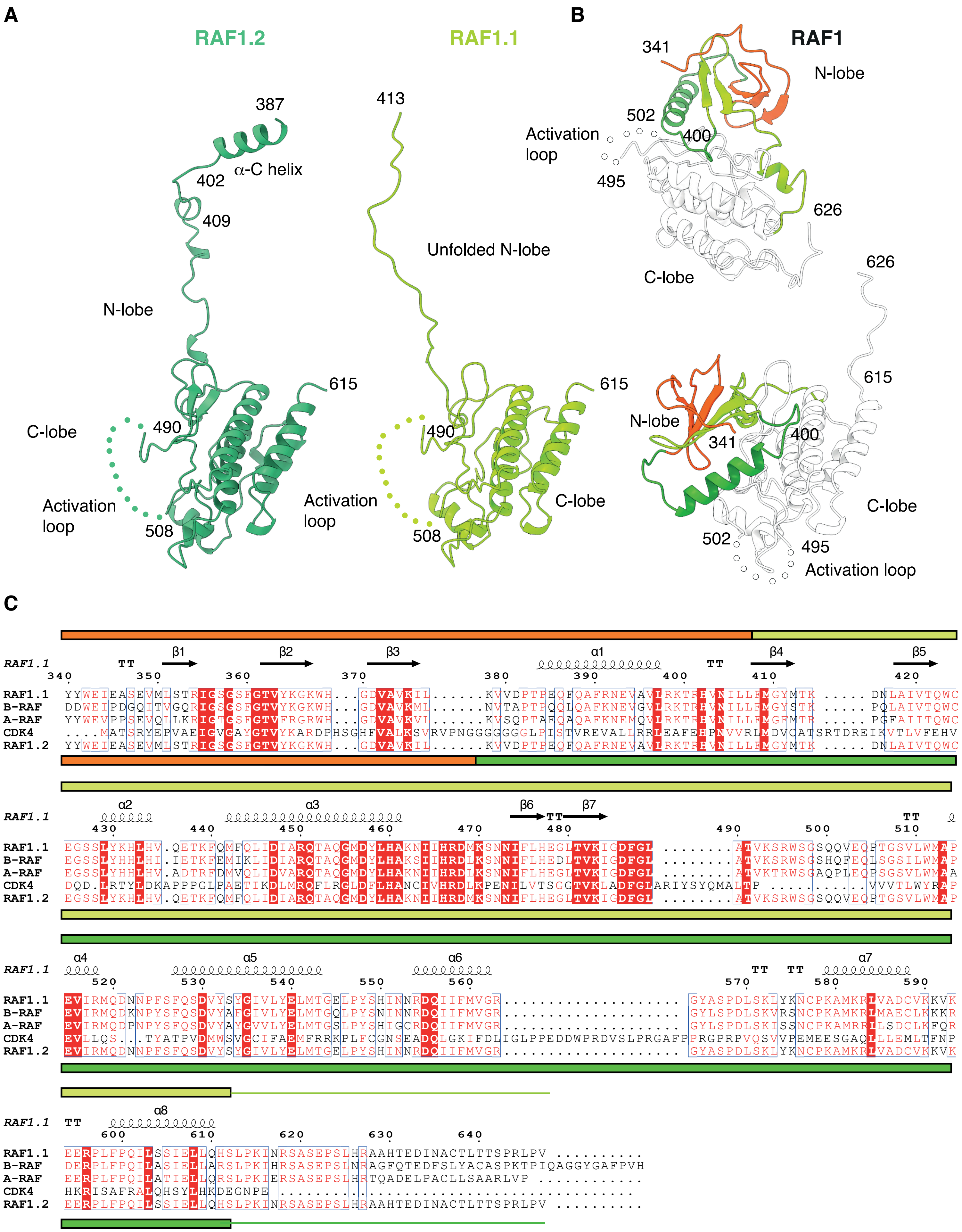

### Fig. S7

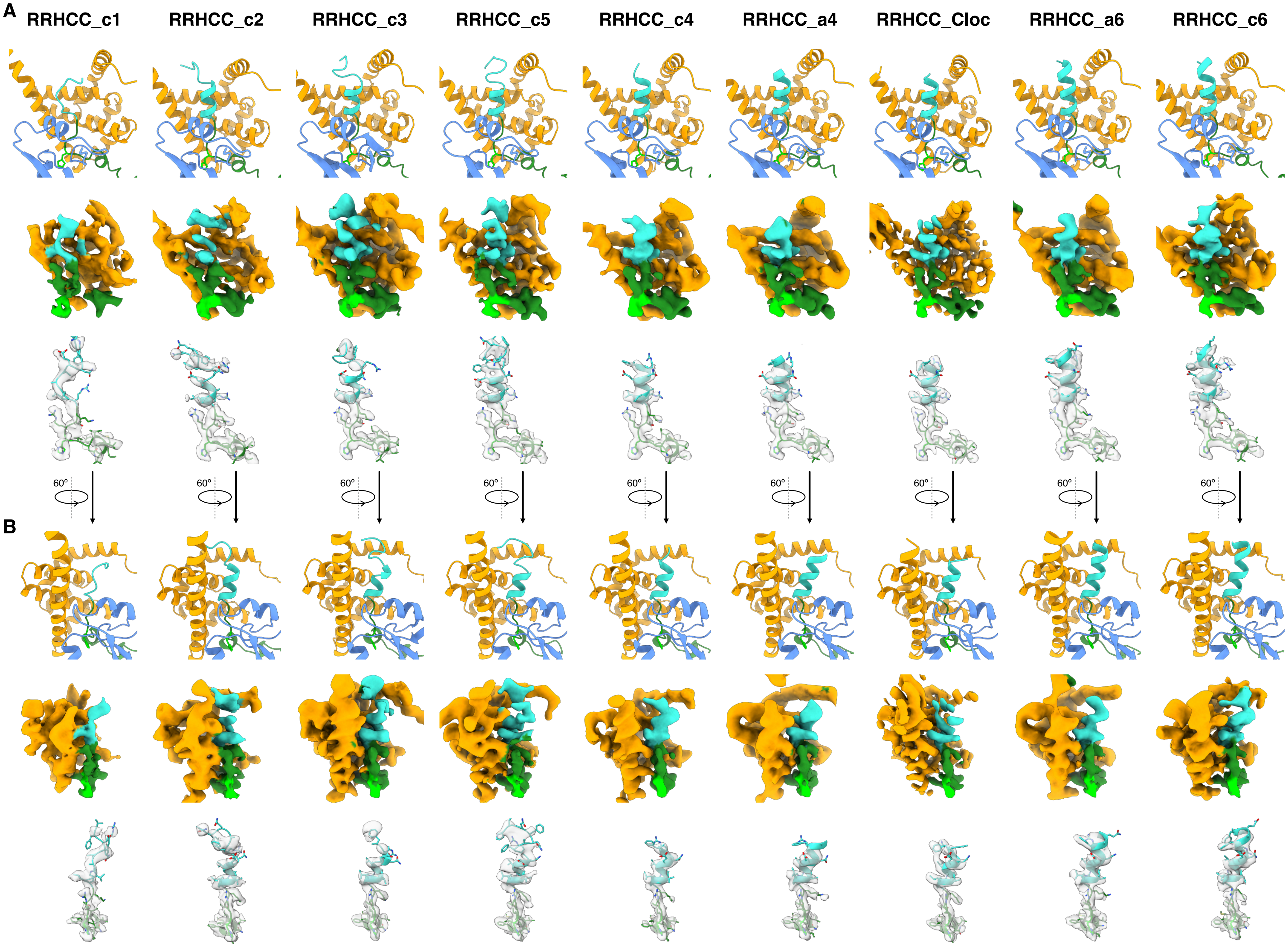

### Fig. S8

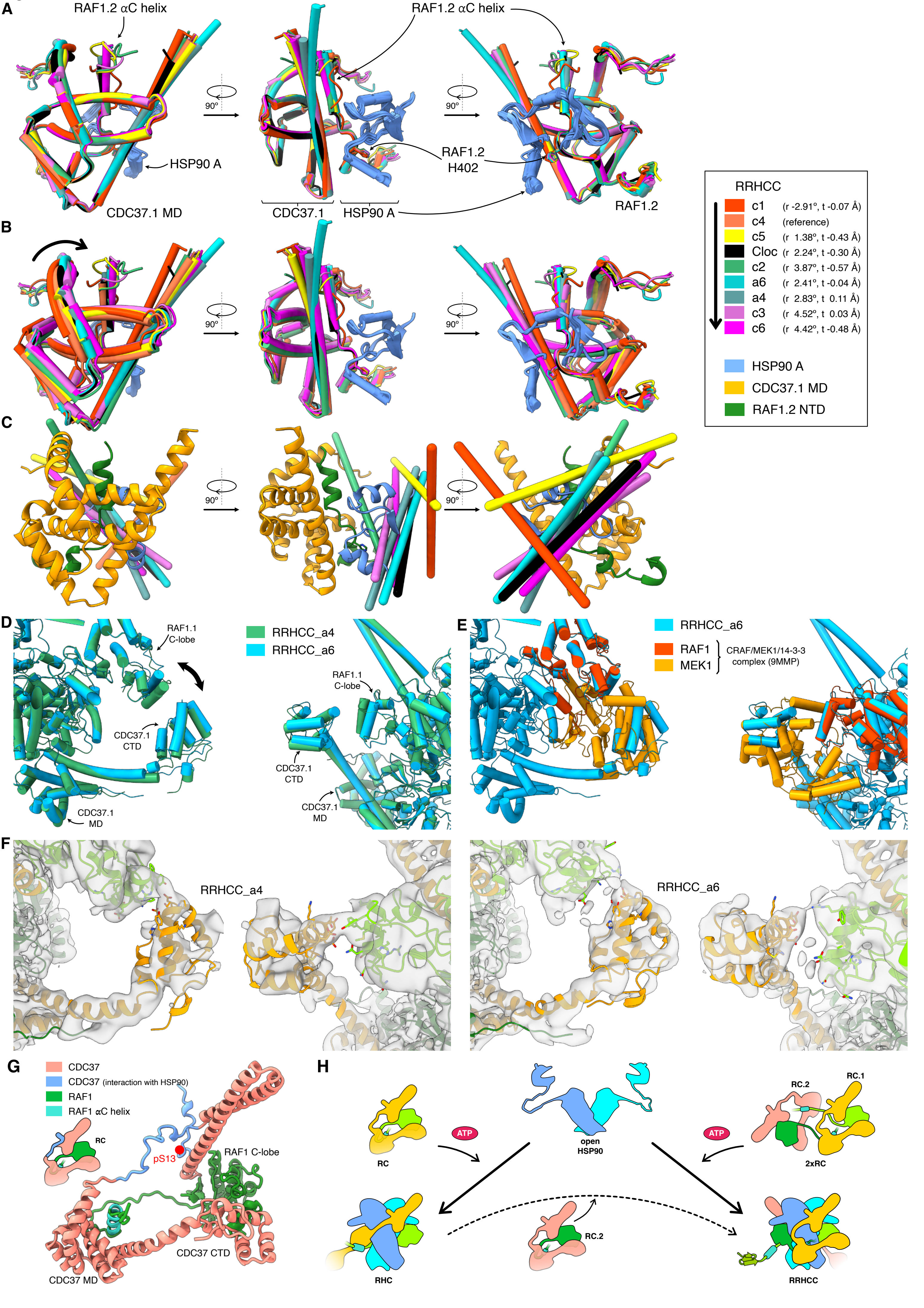

### Fig. S9

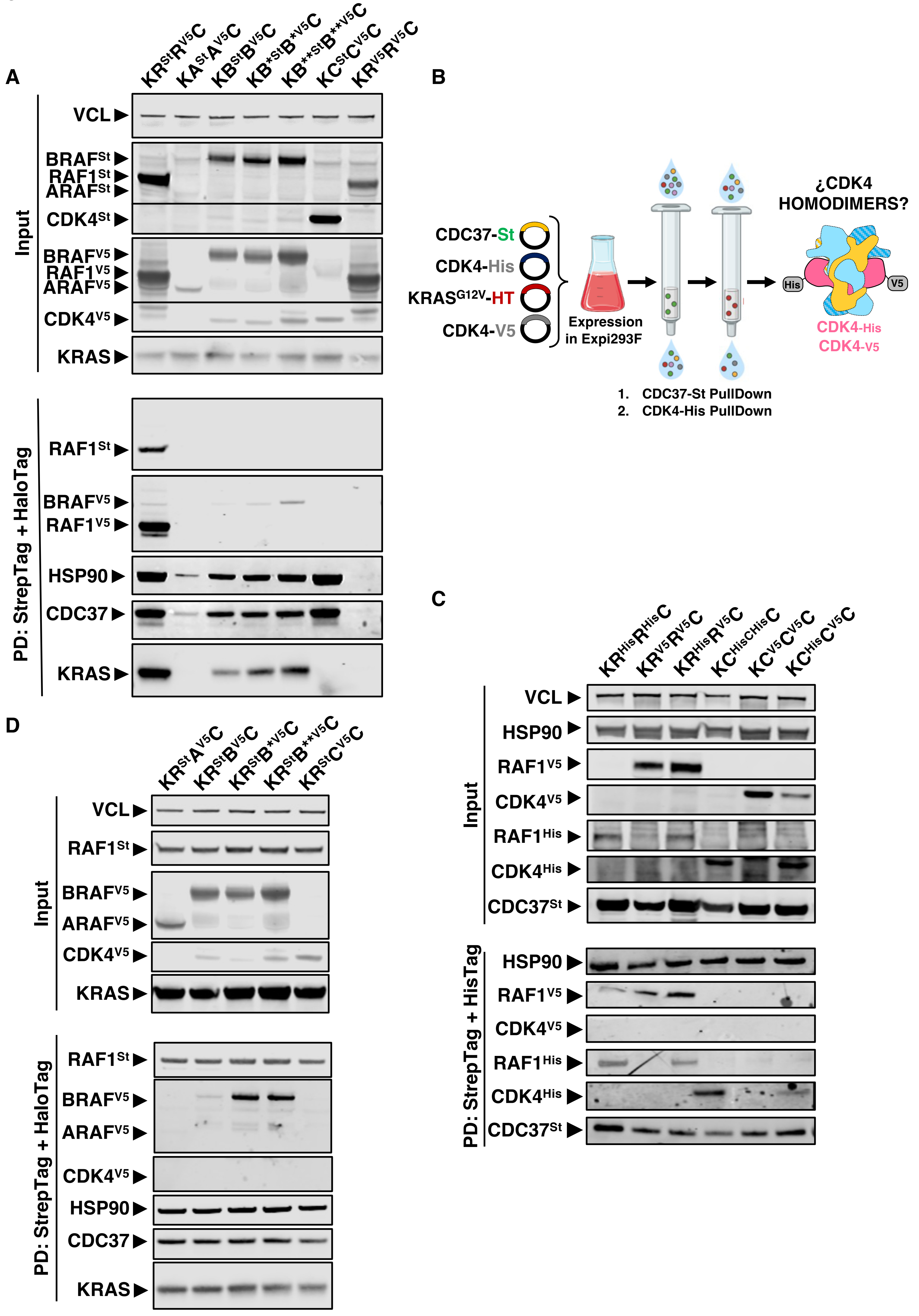

### Fig. S10

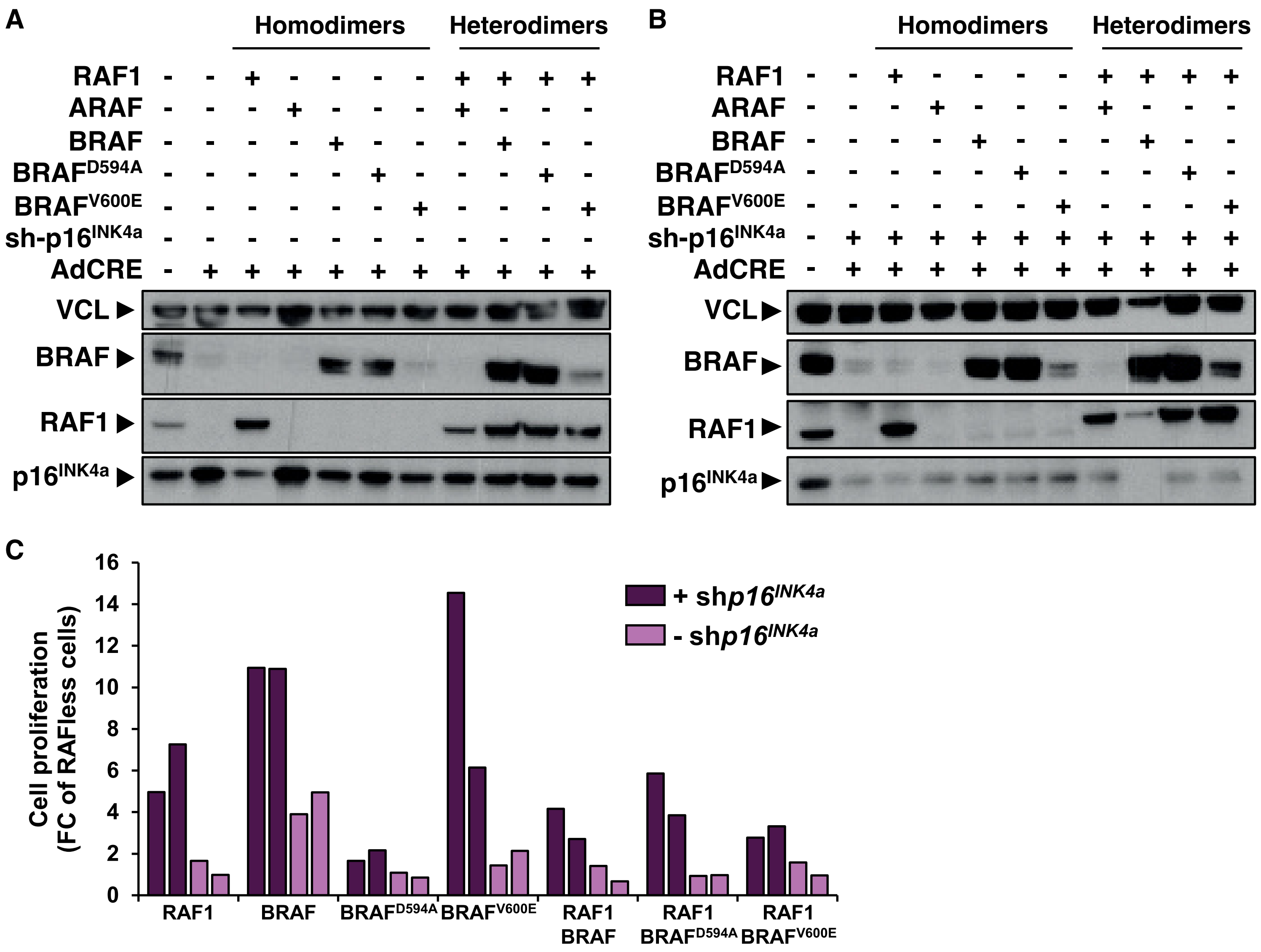

### Fig. S11

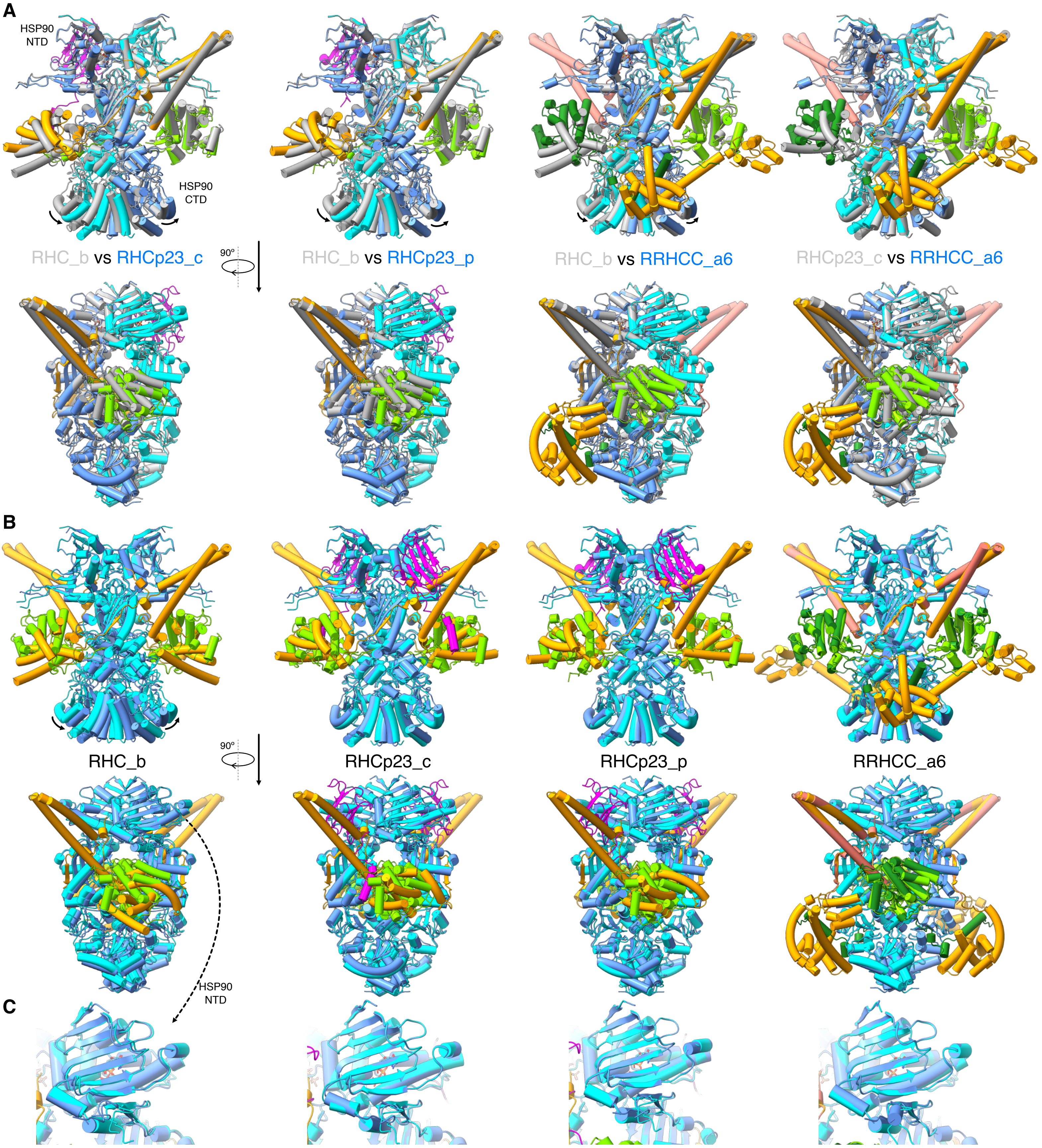

### SI 1

#### Supporting Information 1

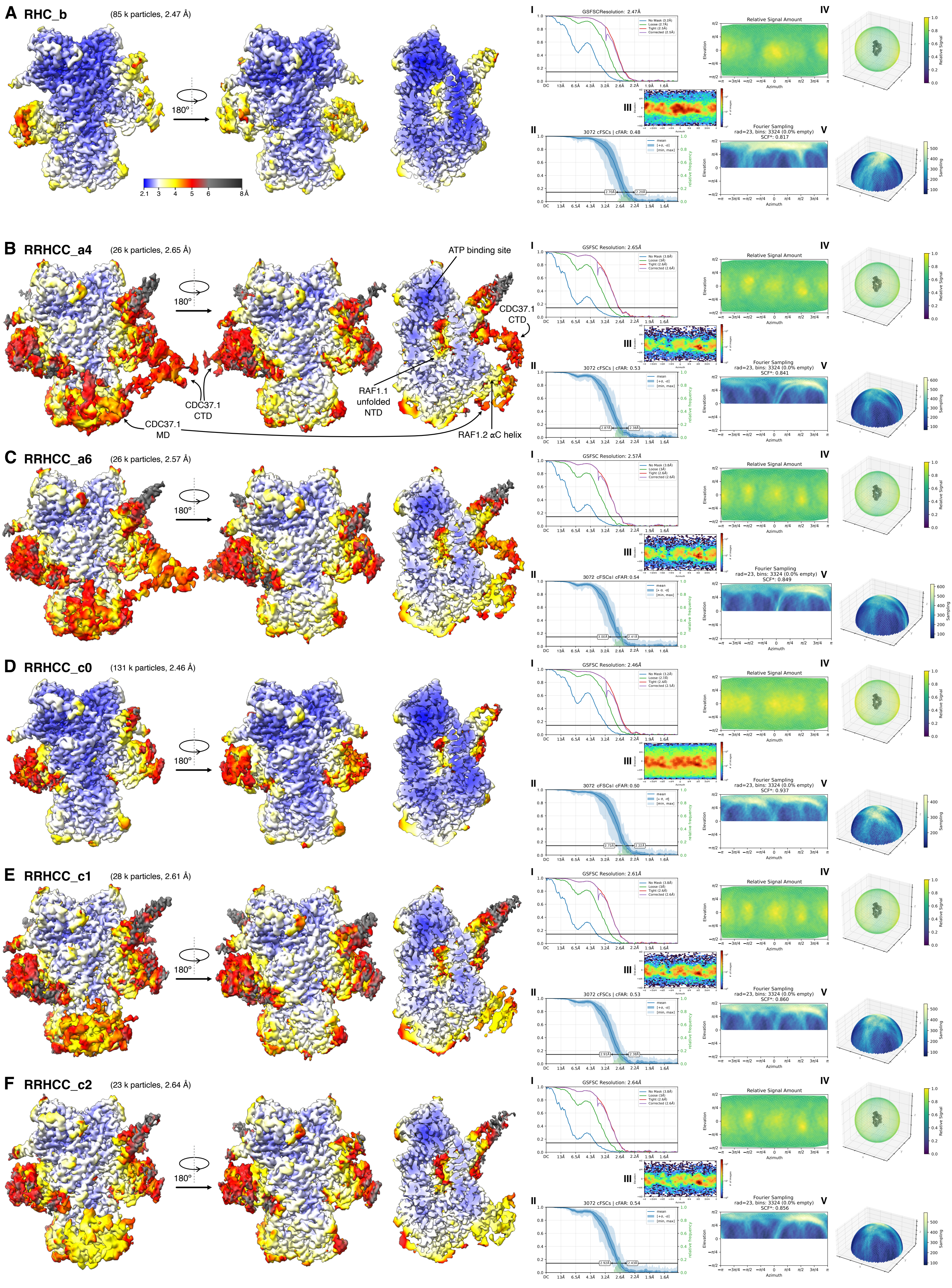

### SI 2

#### Supporting Information 2

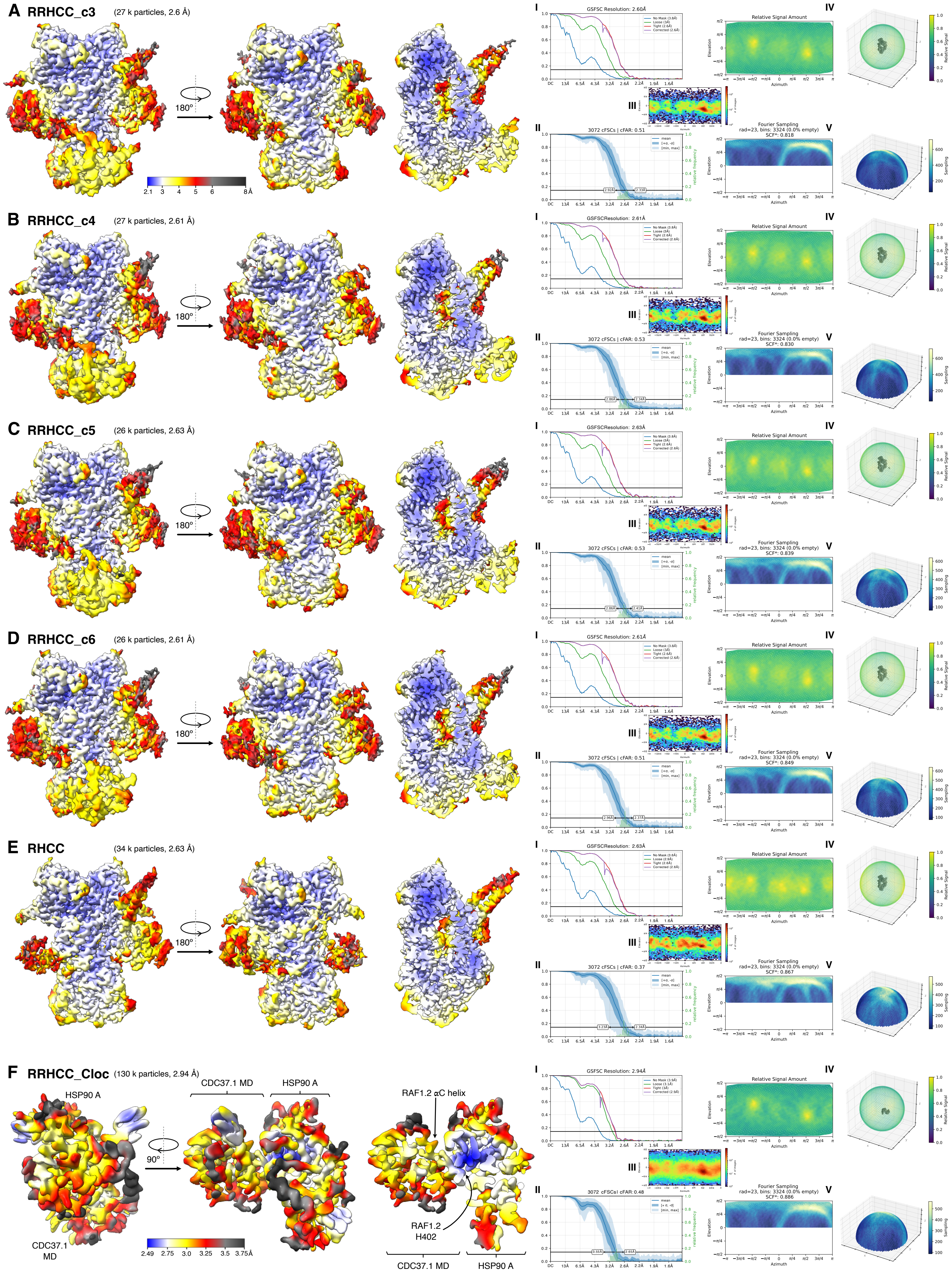

### SI 3

#### Supporting Information 3

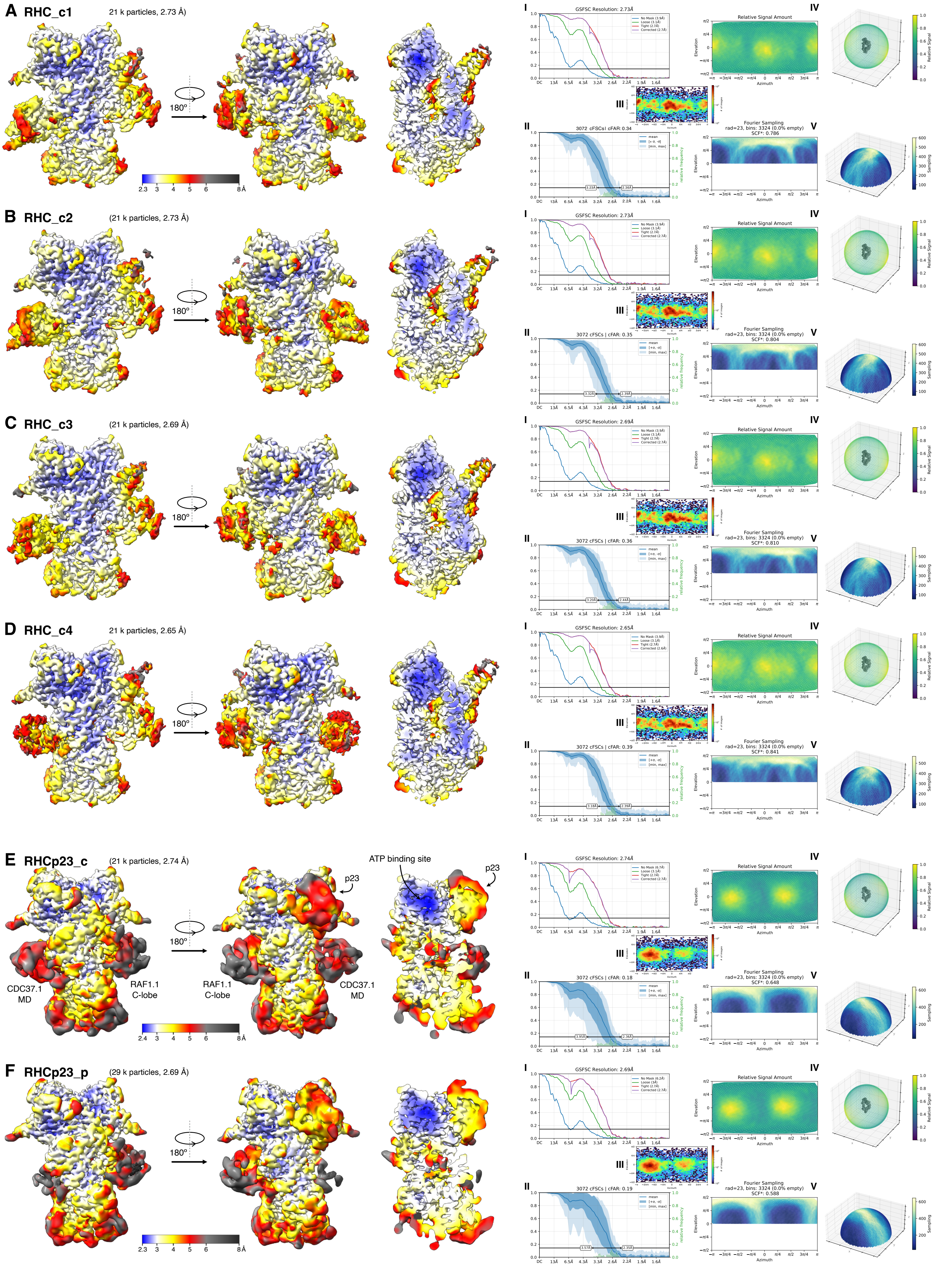

### SI 4

**Supporting Information 4**

A

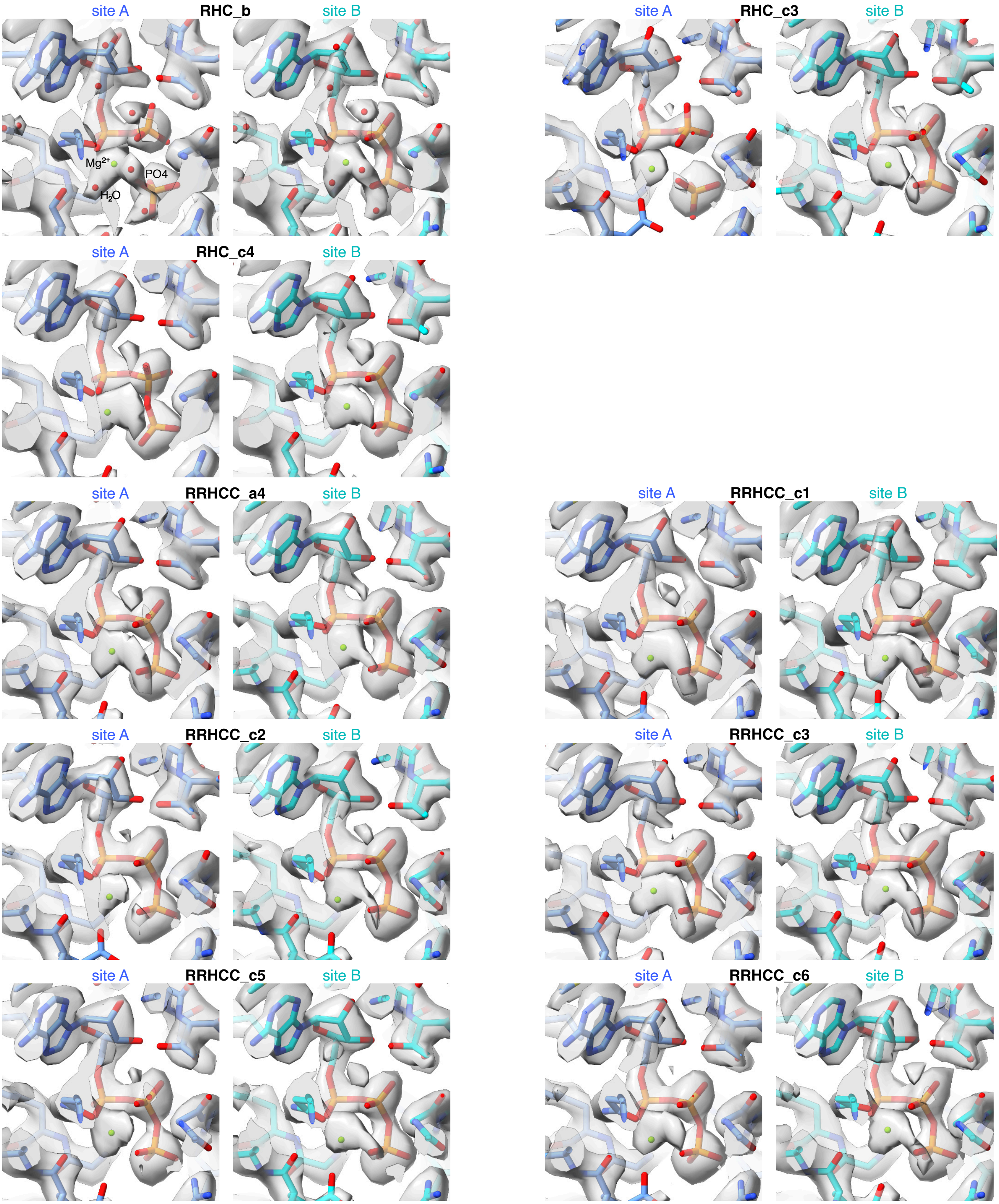


B

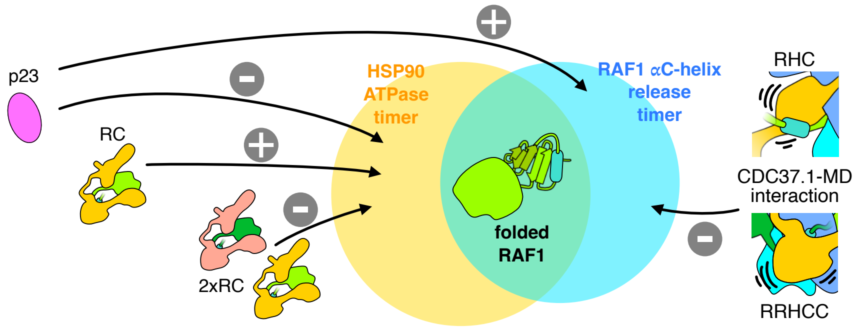
